# Learning regular cross-trial shifts of the target location in serial search involves awareness – an eye-tracking study

**DOI:** 10.1101/2023.12.15.571821

**Authors:** Hao Yu, Fredrik Allenmark, Hermann J. Müller, Zhuanghua Shi

**Affiliations:** General and Experimental Psychology, Department of Psychology, LMU Munich, Munich, Germany

**Keywords:** probability cueing, statistical learning, parallel/serial search, search guidance, eye movements, oculomotor scanning, inter-trial priming, conscious awareness

## Abstract

People can learn, and utilize, not only static but also dynamic (cross-trial) regularities in the positioning of *target* items in parallel, ‘pop-out’ visual search. However, while static target-location learning also works in serial search, acquiring dynamic regularities seems prevented by the demands imposed by item-by-item scanning. Also, questions have been raised regarding a role of explicit awareness for utilizing (at least) dynamic regularities to optimize performance. The present study re-investigated whether dynamic regularities may be learned in serial search when regular shifts of the target location occur frequently, and whether such learning would correlate with awareness of the dynamic rule. To this end, we adopted the same regularity used by Yu et al. (2023) to demonstrate dynamic learning in parallel search: a cross-trial shift of the target location in, e.g., clockwise direction within a circular array in 80% of the trials, which was compared to irregular shifts in the opposite (e.g., counterclockwise; 10%) or some other, random direction (10%). The results showed that ⅔ of participants learned the dynamic regularity, with their performance gains correlating with awareness: the more accurately they estimated how likely the target shifted in the frequent direction, the greater their gains. Importantly, part of the gains accrued already early during search: a large proportion of the very first and short-latency eye movements was directed to the predicted location, whether or not the target appeared there. We discuss whether this rule-driven behavior is causally mediated by conscious control. (248 words).

---

Our visual environment is exceedingly rich and complex, yet our capacity to process information is limited. To make effective use of our cognitive resources, the brain prioritizes information relevant to the task at hand and suppresses irrelevant information that might impede performance (e.g., Treisman and Gelade 1980; Folk, Remington, and Johnston 1992; Wolfe, Cave, and Franzel 1989; Egeth and Yantis 1997). Selection of relevant (and de-selection of irrelevant) information is aided by the structured nature of our environment, which allows us to extract and learn recurrent patterns and regularities that can be beneficial in similar future situations. For example, when looking for our keys, we often commence searching at the ‘usual’ places, like the hallway table or the kitchen counter. Utilizing environmental regularities such as the likely location of a ‘target’ object enables attention and cognitive resources to be deployed efficiently. Effects such as this, referred to as (spatial) ‘probability cueing’, have recently been extensively investigated in laboratory settings. When a task-relevant target appears at a likely location, the attentional system can acquire this information to enhance search efficiency, expediting target detection and attendant response decisions (Druker and Anderson 2010; Geng and Behrmann 2002, 2005; Hoffmann and Kunde 1999; Jiang, Swallow, and Rosenbaum 2013; Shaw and Shaw 1977). Probability cueing is also evident in oculomotor scanning, in terms of an increased frequency and reduced latencies of (early) saccades directed to targets at likely locations (Walthew and Gilchrist 2006; Jones and Kaschak 2012; Jiang, Won, and Swallow 2014). Recently, there have also been demonstrations of an analogous effect: observers may also learn to attentionally suppress the likely location(s) of salient but task-irrelevant ‘distractor’ items in the search displays – referred to as ‘distractor-location probability cueing’ (e.g., Goschy et al. 2014; Sauter et al. 2018; Allenmark et al. 2019; van Moorselaar, Daneshtalab, and Slagter 2021).

It is noteworthy that the majority of studies examining spatial statistical learning (whether of target or distractor locations) did employ some static uneven probability manipulation, such as one display location or region being more likely to contain the target, or a distractor, than any other location or region (e.g., Geng and Behrmann 2002, 2005; Shaw and Shaw 1977; Goschy et al. 2014; Sauter et al. 2018). The probability-cueing effects thus demonstrated were attributed to statistical learning enhancing or suppressing specific (static) locations on the attentional priority map that governs the allocation of focal-selective attention (for a review, see Luck et al. 2021).

More recently, there have been several attempts to extend the study of probability cueing from scenarios with static target and, respectively, distractor distributions to dynamic scenarios, the question being: would statistical learning of selection priorities also work with ‘regular’ – that is, predictable – *changes* in the likely locations of targets or distractors across trials (Li and Theeuwes 2020; Li, Bogaerts, and Theeuwes 2022; Yu et al. 2023). Together, these studies showed that attentional selection can successfully adapt to such dynamic, cross-trial regularities in the placement of *target* items: reaction times (RTs) were faster to targets appearing at the location predicted by the dynamic rule, compared to random locations (Li and Theeuwes 2020; Yu et al. 2023). Importantly, though, Li et al. (2022) found this to critically depend on search being spatially parallel, operating simultaneously across all display items (their Experiment 2b) – rather than serial, proceeding in an item-by-item manner (their Experiment 1), in which case no dynamic cueing effect emerged. In more detail, in Li and Theeuwes’s (2020) design, certain target locations were predictably coupled across trials, such that, for instance, a target occurring at the left-most (or, respectively, the top-most) location in a circular display array on trial *n* would invariably lead to the next target, on trial *n+1*, occurring at the right-most (or, respectively, the bottom-most) location (but not vice versa). When the target was a bottom-up salient shape-singleton item (among differently but homogeneously shaped non-target items), summoning focal attention automatically, participants were able to extract the dynamic target-to-target location shift across trials, as evidenced by facilitated responding to targets at the new, predictable location (compared to targets at non-predictable, random locations). This is consistent with our own study (Yu et al. 2023), in which the search target could also be detected and localized ‘in parallel’; we, too, found that search was facilitated when the target moved predictably across consecutive trials to the neighboring position in either clockwise or anticlockwise (blocked) direction – a somewhat simpler dynamic regularity compared to that introduced by Li and Theeuwes (2020).^1^

In contrast to their own and our parallel search condition, Li and Theeuwes (2020) observed no RT facilitation when the task required search for a (rotated) T-shape target presented among, in terms of feature composition, similar (rotated) L-shaped non-targets – a non-finding replicated by Li et al. (2022)^2^. This task is known to offer little bottom-up or top-down guidance (e.g., Moran et al. 2013), requiring serial scanning of the search array by focal attention to find and respond to the target item. The findings of Li and colleagues (Li and Theeuwes 2020; Li, Bogaerts, and Theeuwes 2022) would then suggest that dynamic, cross-trial regularities in the placement of the target item may not be extractable and used to improve performance under conditions of serial search.

Thus, with static (spatially fixed) likely target locations, target-location probability learning works under serial as well as parallel search conditions (Geng and Behrmann 2002). With dynamic target-location regularities, by contrast, it appears to work only under parallel, but not serial, conditions (Li and Theeuwes 2020; Li, Bogaerts, and Theeuwes 2022). The question is: why?

## Why would *dynamic* target-location probability-cueing be dependent on the – parallel vs. serial – search mode?

While Li and colleagues offer little by way of an explanation, a possible answer may have to do with the complexity of monitoring attention allocations over time, within and across trials. Under conditions of parallel search, the target ‘pops out’, that is, it is nearly always the first and, in fact, the only item that summons attention – the only item because, when the target is selected, it is identified as the task-relevant item and the response-critical information can be extracted and search terminated. As a result, the current target location is ‘marked’ by the system as being task-critical, and this may allow some higher-order ‘working-memory’ system that monitors attention allocations over time (where was attention allocated to and where is it to go next?) to pick up cross-trial dependencies in the positioning of consecutive targets within a regularly structured (circular) display array.

Under serial conditions, by contrast, search involves attentional inspection of a (varying) number of non-target items before the target is eventually selected, upon which the search is terminated. Monitoring attention allocations over time is considerably more complex, as the locations of already inspected non-target items would need to be marked and held in some type of memory to prevent re-visitations. As a result, the location of the target, once eventually selected on a given trial, would stand out less, compared to the location of a pop-out target. In addition, the search on the next trial might again start with some more or less randomly selected (likely a non-target) location, which would make it more difficult to track dynamic regularities of the target placement across trials.

Compared to dynamic conditions in which the target location changes regularly across trials (i.e., the target location is [almost] never the same on the current as on the preceding trial), static regularities would be easier to pick up even under serial search conditions, as scanning would almost always end up at exactly the same location, allowing knowledge of the fixed target-location probabilities to be gradually accumulated across sequential trial episodes.

Thus, the increased online-, or working-, memory demands in monitoring attention allocations (within trials) and search-terminating target locations (across trials) under serial vs. parallel search would particularly impact the acquisition of dynamic regularities in the target placement. In contrast, static regularities may be extracted relatively efficiently even in serial search. Nevertheless, we hypothesize – and test in the present study – that, depending on the frequency with which a dynamic rule is invoked as well as possibly its complexity, participants may be able to extract the regularity even in serial search and use it to optimize performance.

### Is (dynamic) target-location probability cueing implicit in nature?

It is widely assumed that statistical learning is implicit in nature, extracting statistical regularities from the input without explicit awareness or intent (Turk-Browne, Jungé, and Scholl 2005; Turk-Browne et al. 2009). Consistent with this are reports that individuals can learn and utilize static regularities related to the locations of salient distractors without awareness; that is, most participants were unable, in post-experimental awareness tests, to point out the frequent distractor location, and the cueing effect differed little between those who correctly selected vs. those who failed to select the frequent location (e.g., Failing, Wang, and Theeuwes 2019; van Moorselaar and Theeuwes 2022; Failing et al. 2019; Wang and Theeuwes 2018). There have been similar findings with regard to the statistical learning of target locations (e.g., Li, Bogaerts, and Theeuwes 2022; Geng and Behrmann 2005; Ferrante et al. 2018).

However, the idea that probability cueing is implicit in nature has come under scrutiny with studies using more sophisticated awareness measures to probe the relationship between explicit awareness and the cueing of target locations (van Moorselaar and Theeuwes 2023; Huang, Donk, and Theeuwes 2022; Yu et al. 2023; Golan and Lamy 2023; Giménez-Fernández et al. 2020; Vicente-Conesa et al. 2021). The conflicting indications regarding a role of awareness in the learning of static target regularities may arise from a variety of factors, such as the probability levels used in the various studies, the number of trials, and the method employed to assess awareness (Jan Theeuwes, Bogaerts, and van Moorselaar 2022). For instance, asking participants to rank the possible locations from the most probable to least probable and estimate the number of times that the target had appeared in each display quadrant (in a “serial”, contextual-cueing paradigm; cf. Chun and Jiang 1998), Giménez-Fernández et al. (2020) found that many participants were actually aware of the target’s unequal (*static*) spatial distribution. In a recent study of *dynamic* target-location probability cueing in pop-out search (Yu et al. 2023), we likewise found a substantial number of participants to be explicitly aware of the dynamic (cross-trial) target regularity, and we observed the dynamic target-location probability-cueing effect to be significant only in the group of aware participants.

Based on these findings, we hypothesize that at least the learning of *dynamic* target-location regularities in *serial* search is explicit in nature, dependent on (or correlated with) participants becoming aware of the rule governing the shifts in the target location across trials.^3^

### Role of inter-trial target-identity swapping, positional priming, and rule-based priming

Besides serial search making greater demands on the tracking of attention allocations (within trials) and target placements (across trials), the difficulty can be further heightened if the target identities (e.g., shape) change randomly, alternating with the non-target’s identities across trials, as opposed to remaining fixed. Note that feature swapping is a standard feature in ‘additional-singleton’ (i.e., pop-out target plus distractors) paradigms (e.g., Jan Theeuwes 1991), where it is designed to promote a spatially parallel ‘singleton-detection’ search mode (cf. Bacon and Egeth 1994). In such paradigms, statistical learning of distractor locations is influenced by whether or not there is random feature swapping across trials (e.g., Allenmark et al. 2019; Zhang et al. 2019), likely because further processing is required to establish the (dimensional or featural) identity of both the distractor and target items. Of note, swapping of the color that singled out the target from the color-homogeneous background items was also implemented in Yu et al.’s (2023) dynamic target-location learning experiment. This did not hinder (aware) participants from acquiring the dynamic rule, likely because the target popped out of the search array.

Random swapping of target and non-target features is less common in studies of serial search. Conceptually, without swapping, observers can set up a fixed ‘target template’ for comparing any selected item to make a target/non-target decision. This also potentially allows for a top-down biasing selection towards critical features that differentiate the target from non-target items. In contrast, with swapping, observers would need to create two templates and determine, for each trial, which one is the target template and which the non-target template. Establishing this would require inspection of multiple items: if two inspected items share essentially the same features, they are likely non-targets – yielding the definition of the non-target template. So, by default, the other description becomes the target template. Typically, under conditions of swapping, the search system carries over the template from one trial to the next (e.g., Maljkovic and Nakayama 1994; Kristjánsson, Wang, and Nakayama 2002; Geyer, Müller, and Krummenacher 2006) – the implicit assumption being that critical task settings stay the same and additional information is required to change, or update, the task set, expediting search on no-swap (vs. swap) trials. Nevertheless, given the added complexity, in terms of required attention allocations, to establish the identity of the target (template) under conditions of random swapping, one would expect dynamic target-location learning to be less robust under conditions of randomly variable vs. fixed target identity.

Note that two other types of intertrial priming may be at work, especially under conditions of serial search. The first is *positional intertrial priming* (e.g., Maljkovic and Nakayama 1996; Krummenacher et al. 2009), characterized by the attentional priority being raised for the target location on a given trial and carry-over of this positional selection bias to the next trial. This type of intertrial priming might be particularly prominent under (serial) search conditions that provide no other (e.g., feature-based) sources of guidance to the target location. In this situation, the system might strongly prioritize inspection of locations where a target was detected in the previous search episode. Any dynamic rule-based target-location probability cueing effect would have to compete with this positional priming effect, which thus provides an important reference against which to compare the probability cueing effect.

Finally, assuming that a dynamic target-location regularity is acquired, in the form of some kind of top-down ‘prior’ predicting the next location, the weight of this prior on a given trial might depend on whether the target placement on the preceding trial was consistent with the rule (rule-conforming) or inconsistent (rule-breaking). Rule-conforming target placements might strengthen (the weight of) the rule, whereas rule-breaking placements might weaken the rule – yielding a *rule-based intertrial-priming* effect. Again, rule-based priming effects might be particularly prominent under (serial) search conditions that offer no, or few, other sources of guidance to the target location.

### Objective and rationale of the present study

The present study aimed to examine whether participants would learn a simple dynamic (probabilistic) regularity in the positioning of target items across consecutive trials in a *serial* search task, and if such dynamic learning would rely on explicit awareness of the regularity. We used the same dynamic, cross-trial regularity as Yu et al. (2023) did in a parallel search task. This involved shifting the target location in a circular display arrangement by one position, either clockwise or anticlockwise (blocked per participant) across trials with a probability of 80% (see Figure 1 for a depiction of the search displays and the dynamic regularity in the positioning of sequential target items).

**Figure 1.**
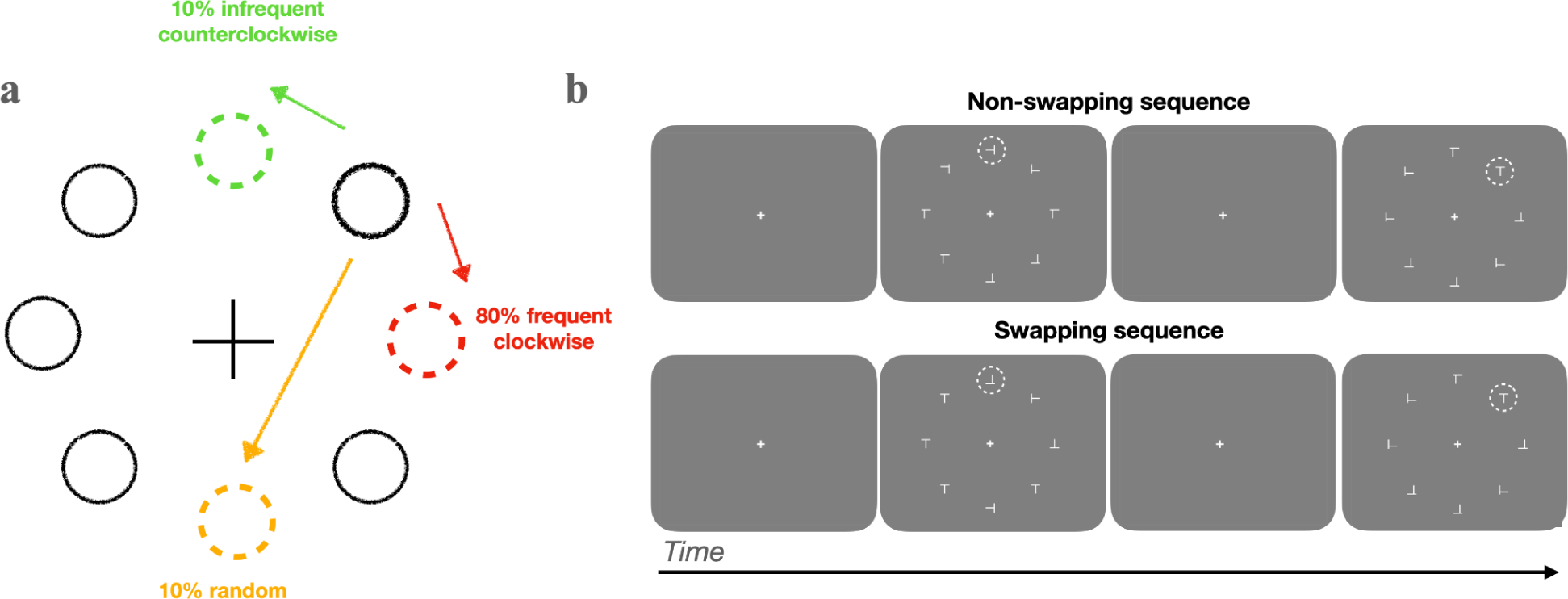
(a) Illustration of the three cross-trial target-location transition conditions. There were three types of change of the target location across consecutive trials: With 80% probability, the critical item would move to the adjacent location, in either clockwise or counterclockwise direction (here, indicated by the red dashed circle marking the ‘frequent’ location). The frequent direction was fixed for a given participant and counterbalanced across participants. With 10% probability, the critical item would shift to the adjacent location in the opposite direction (indicated by the green dashed circle marking the ‘infrequent’ location). In the remaining 10% of trials, the critical item would move randomly to any of the other locations, including re-appearing at the same location (indicated by the yellow dashed circle marking a ‘random’ location). (b) Examples of sequences from trial blocks with random swapping (mixed) and, respectively, no-swapping (fixed) of the target identity across trials. In the mixed condition, the target identity changes randomly from trial to trial; in the fixed condition, it stays the same.

With the regular shifts occurring in 80% of the trials, compared to only a 25%-probability in Li and Theeuwes (2020), we expected a substantial number of participants would extract and utilize this regularity to speed performance even in serial search, which requires (a sequence of) eye movements to detect and respond to the target item. In particular, we expected faster task-final RTs if the cross-trial shift of the target location conforms with the rule (‘frequent’-shift trials in Figure 1) versus when it didn’t (‘infrequent’- and ‘random’-shift trials), evidencing a dynamic target-location probability-cueing effect.

Inspired by the findings by Yu et al. (2023), we expected that only those participants who, based on a post-experiment awareness test, were ‘aware’ of the dynamic regularity, would exhibit a dynamic target-location probability-cueing effect. ‘Unaware’ participants, by contrast, were not expected to show a benefit from the regularity. We also expected a correlation between the subjective certainty about the rule and their cueing effect.

In addition to examining the search-final RTs, we also tracked participants’ eye movements while they scanned through the search displays for the target. RTs reflect the culmination of various processes contributing to the final response decision timing, and as such (e.g., without sophisticated methods to decompose RTs) they are limited in revealing which component processes came into play at what time during a trial in producing the response required by the task. In this regard, the tracking of eye movements provides critical data – in particular, in complex search tasks requiring the serial allocation of attention, which inherently involves sequential eye movements to find and respond to the target item. Accordingly, here, we examined participants’ eye movement in order to gain further insights into the time course of *dynamic target*-location probability cueing (for oculomotor studies of static distractor-location probability cueing, see, e.g., Allenmark, Shi, et al. 2021; Di Caro, Theeuwes, and Della Libera 2019; Sauter et al. 2021; B. Wang, Samara, and Theeuwes 2019). In fact, our task required participants to expressly fixate the target item and, upon confirming it as the target, execute a simple manual detection response.

Thus, recording participants’ eye movements allowed us to examine, in aware participants, at what stage(s) of the search saccadic behavior would be guided by the acquired rule – or regularity ‘prior’ –, over and above any bottom-up and top-down guidance signals afforded by the search task as such. In particular, if rule-based guidance kicks in very early, already the first saccade (from the initial, central fixation spot) might tend to be directed straight to the dynamically predicted, ‘frequent’ target location, compared to other locations – in particular, an ‘infrequent’ position in the opposite direction to the (clockwise/anticlockwise) rule that shares the same distance from the last target location as the ‘frequent’ location, or compared to the same location occupied by the target on the last trial (positional intertrial priming). In any case, even if rule-based guidance takes longer (than the first eye movement) to come into play, we would expect that aware participants would use fewer saccades to locate the target at the frequent location compared to other locations (except possibly the repeated one), and fewer than unaware participants. This oculomotor dynamics would eventually manifest in cueing effects in the search-final RTs.

Additionally, by mixing ‘frequent’ and ‘random’ (baseline) target placements within blocks, rather than segregating them into separate blocks (e.g., L. Wang, Wang, and Theeuwes 2021), we could assess how dynamic rule guidance on a given trial is modulated by preceding trial events conforming that either conform to or break the rule (rule-based intertrial priming). The eye-movement record can trace this influence back to even the earliest saccades executed on a trial.

Relatedly, we examined these issues under conditions where the target (vs. non-target) identity remained constant across trials and, respectively, under conditions where target and non-target identities were mixed, swapping randomly across trials. The additional task demands imposed by the latter condition would make the task harder, forcing extended serial scanning (of several items) to determine the target and non-targets on each trial. As just inspecting the item at the location predicted by the dynamic rule would not be sufficient to confirm its target status, the mixed condition may weaken (or interfere with) rule application or, conversely, strengthen reliance on the rule, as all relevant information for decision-making would likely be available at the predicted location (the frequent target position) and its vicinity (likely containing a non-target item). Again, early eye movements would provide insights into the (sub-) processes generating the task-final RTs under these conditions.

Finally, in addition to examining whether any probability-cueing effects in the task-final RTs correlate with participants’ awareness of the dynamic regularity, recording eye movements allows us to examine whether already the earlies saccades executed during serial search may be ‘informed’ by explicit knowledge where the new target is likely to be located.^4^

## Method

### Transparency and Openness Statement

Our report details the methodology used to determine the sample size, incorporating both a theoretical comparison and a power analysis. We also fully disclose the criteria for data inclusion and exclusion in pre-processing and all subsequent analyses. Regarding these criteria: no participants were excluded from the study, and all criteria for trial-based inclusion and exclusion were pre-determined prior to data analysis. We report all data manipulations in the study. The experimental code, raw data, and data analyses of the present study are publicly available at: https://github.com/msenselab/learning-in-serial-search. The experiment was conducted in 2022.

### Participants

24 healthy university students from LMU Munich participated in this study (mean age ± SD: 26.9 ± 4.1 years; ranging from 20 to 33 years; 21 females, 3 males). All participants reported normal or corrected-to-normal vision, and passed the Ishihara color test (Clark 1924), confirming unimpaired red-green color perception.

To ensure robust statistical power for addressing the questions at issue, we estimated our sample size based on our previous study (Li and Theeuwes 2020; Li, Bogaerts, and Theeuwes 2022; Yu et al. 2023), which employed a similar manipulation of the dynamic (cross-trial) target-location regularity and reported an effect size of *f* = 0.42 (average across all experiments). An a-priori power analysis, conducted with an effect size of *f* = 0.42, an α = .05, and 98% power (1–β), indicated a minimum sample size of *n* = 20 (G*Power 3.1; Faul et al. 2007). Given that our study introduced a more complex letter-search paradigm, we increased the sample size to 24.

The study was approved by the LMU Faculty of Pedagogics & Psychology Ethics Board. All participants provided written informed consent prior to the experiment and received 9.00 Euro per hour or equivalent course credits for their participation.

### Apparatus

The experiment was conducted in a sound-attenuated, dimly lit testing chamber. Participants were seated 55 cm away from a 24-inch CRT display monitor that displayed the search stimuli at a screen resolution of 1920 × 1080 pixels and a refresh rate of 120 Hz. We employed PsychoPy (v. 2022.2.2) to control stimulus presentation, manual-response recording, and eye-movement tracking.

Gaze position for the dominant eye was captured using an SR Research EyeLink 1000 desktop-mount eye-tracker (Osgoode, Ontario, Canada), operating at a sampling rate of 1kHz. Participants registered their responses using a QWERTZ keyboard by pressing the space button with either their left- or right-hand index finger.

### Stimuli and Design

The search displays (see Figure 1) featured a white fixation cross at the center, set against a gray screen background. Each display contained eight items: a single target shape, either a “T”- or “L”-shaped letter, among seven non-target shapes, “L”- or “T”-shape letters). When the target was a “T”, the non-targets were all “L”-shaped, and vice versa.

The eight display items, each subtending 1.25° × 1.25° of visual angle (CIE [Yxy]: 70.5, 0.330, 0.326), were equal spaced on a virtual circle, at an eccentricity of 7° (yielding a center-to-center distance of 5.4° between adjacent items). To elevate task difficulty and encourage serial search, the “L”-shaped items featured a slight offset at the line junction, measuring 0.3°. Both “T” and “L” shapes appeared randomly in one of the four orthogonal orientations (0°, 90°,180°, or 270°). A shape-defined target, either a “T” or an “L”, was present on every trial. The target could appear at any of the eight possible display locations, with its location uniformly distributed across all trials. Participants were tasked to locate the target with their eyes (i.e., making a saccade to it and fixating) and then promptly press the spacebar to confirm target identification. Upon their response, a feedback message showed for 500 ms, indicating either “Correct (response)” in green or “Incorrect (response) ” in red.

Crucially, the positioning of the target within the circular array was probabilistically predictable across consecutive trials *n* and *n+1*. In 80% of the trials, the target shifted to an adjacent location, in a consistent clockwise or counterclockwise direction – we refer to this as the “frequent (target) location”. The primary direction of this shift was constant for each participant, but counterbalanced across participants. In another 10% of the trials, termed “infrequent condition”, the target moved to an adjacent location in the opposite direction to that of the frequent condition. For the remaining 10% – the “random condition” – the target’s position was chosen randomly among the six remaining alternative locations (including repeated presentation at the same location). Note that upon any irregular shift (including “infrequent” shifts one by position in counter-direction, position repetitions, and any larger “random” shifts), a regular shift (in the “frequent” direction) on the subsequent trial would proceed from the last target location. This is exactly the same dynamic regularity introduced in Yu et al.’s (2023) parallel-search Experiment 1.

The experiment consisted of 16 blocks: 8 “target-fixed” blocks, in which the target remained the same across trials, were randomly interleaved with the other 8 “target-swapping” blocks (in which the target identity changed randomly from trial to trial). Each block consisted of 60 trials, yielding a total of 960 trials for the whole experiment. Of note, the target-swapping condition of the target served as a within-subject variable manipulated between blocks. That is, the shape of the target (as well as that of the other, non-target items) could randomly swap across trials, in line with prior studies (of mainly singleton) search (e.g., Allenmark et al. 2019; J. Theeuwes 1992).

### Procedure

Each trial began once a stable fixation on the central fixation cross was detected (i.e., fixation within a virtual circle of 2° radius for at least 500 ms). Following a randomized (fixation) duration between 700 and 1000 ms, the circular search array was presented and remained visible until the participant responded.

Participants were instructed to localize the target within the display array by making an eye-movement to it and then press the spacebar as fast as possible to confirm that they had actually located the target (rather than a non-target item); they were told that they were free to move their eyes in their search for the target. A trial was marked as ‘correct’ when participants fixated on the target item (i.e., within a circular region of 2.5° radius centered on the target) during the key-press response. If participants fixated a non-target item or no item at all, the feedback message “Incorrect” appeared at the screen center for 500 ms. Each new trial started with the reappearance of the central fixation cross. Between blocks, participants could take a break of a self-determined length.

To determine participants’ awareness regarding the dynamic regularity of the target locations across trials, a post-experimental questionnaire was administered. It consisted of three forced-choice questions: First, participants had to indicate whether or not they had noticed *any* regularity in the target’s placement across trials, selecting from six options (Was there any regularity? – Definitely no; Probably no; Possibly no; Possibly yes; Probably yes; Definitely yes). Second, they had to specify the dominant (regular) direction of the movement, by choosing one of two options for the most frequent type of movement (moved clockwise; moved counterclockwise.) Third, based on their previous answers, they estimated the frequency, in percentage terms, of the target moving in that direction (from 0% to 100%).

### Eye-data pre-processing

The recorded eye-position data were analyzed off-line. Saccades were identified based on their velocity distribution, using a moving average over twenty successive eye-position samples (Engbert and Mergenthaler 2006). Default settings were used to determine the on- and offset of saccades. A saccade was marked as landing on the target, or a non-target, if its endpoint fell within 2.5° from the center of the respective item (see Figure 1b). Trials with response errors (i.e., participants pressing the spacebar while fixating outside the target region) were relatively low (4.9%) and excluded from further analysis.

## Results

### Awareness test

Given our recent finding (Yu et al., 2023) that awareness plays a – likely critical – role in the learning of dynamic cross-trial regularities, we first classified participants into an ‘aware’ and an ‘unaware’ group and then examined search performance separately for the two groups. Among the 24 participants, 16 reported having noticed “a regularity” in the cross-trial target movement, and all of them correctly identified the specific type of regularity they had actually encountered in the search displays. These participants were assigned to the aware group. The remaining eight participants, who, according to their answers to the questionnaire, were unable to identify the regularity formed the unaware group.

### Response times

RT analyses were performed on individuals’ mean RTs after excluding error trials (i.e., trials in which participants did not fixate within the 2.5° region around the target but gave a manual, spacebar response, which happened in approximately 4.9% of the trials, on average). Figure 2 depicts the mean RTs (calculated across individual participants’ means) for the three cross-trial Target-Location Transition conditions (frequent, infrequent, random), separately for the two Target-Constancy block types (target identity fixed vs. mixed) and the two groups (aware vs. unaware).

**Figure 2.**
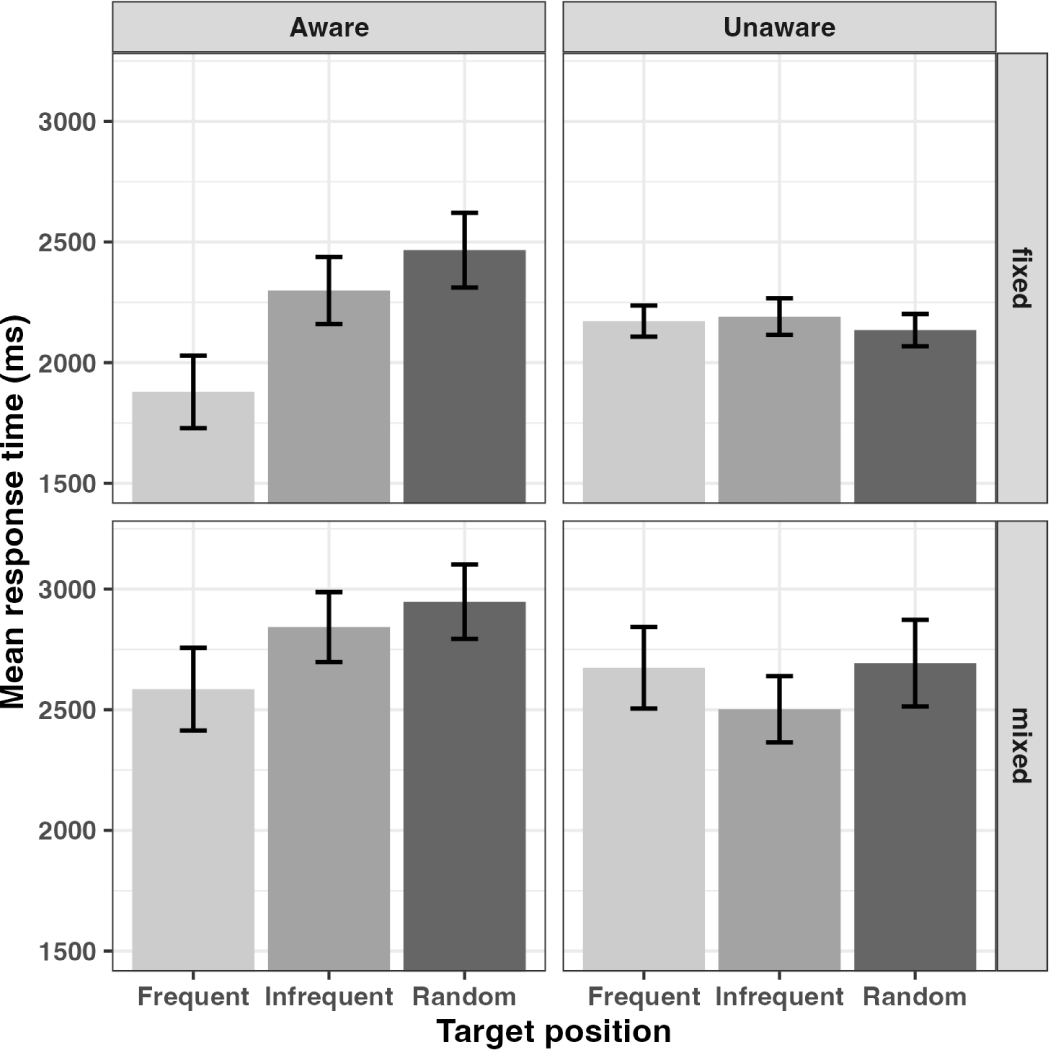
Mean RTs as a function of the cross-trial target-location transition (random, infrequent, frequent transition) and cross-trial target constancy (target identity fixed, mixed per block), separately for the aware and the unaware groups of participants. Error bars represent one standard error.

A mixed ANOVA with the within-participant factors cross-trial Target-Location Transition (random, infrequent, frequent) and cross-trial Target Constancy (target identity fixed, mixed per trial block) and the between-participant factor Awareness (aware, unaware) revealed significant main effects of Target Constancy, *F*(1,22) = 53.266, *p* < .001, 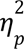 = 0.708, and Location Transition, *F*(2,44) = 5.698, *p* = .006, 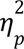 = 0.206. RTs were overall (by > 500 ms) faster when the target identity was fixed per block compared to when it changed randomly across trials; and RTs were overall faster when the target location shifted by one position in the frequent direction across trials (2328 ms) compared to both a shift by one position in the infrequent (i.e., counter-) direction (2459 ms) or a random shift (2560 ms). Additionally, the Location-Transition × Awareness interaction was significant, *F*(2,44) = 7.182, *p* = .002, 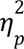 = 0.246 – which was due to only the aware group, but not the unaware group, showing a systematic Location-Transition effect. As revealed by post-host comparisons (with Bonferroni correction), for the aware group, RTs were faster in the frequent transition condition (2232 ms) relative to both the infrequent condition (2571 ms), *t*(15) = 3.534, *p* = .004, *d_z_* = 0.554, and the random condition (2707 ms), *t*(15) = 4.954, *p* < .001, *d_z_* = 0.777, with only a numerical difference between the latter (*t*(15) = 1.421, *p* = .497, *d_z_* = 0.22). In contrast, there was no significant facilitation by the frequent transition for the unaware group (both *p*’s = 1, *d_z_*s < 0.66). This indicates that participants in the aware group (but not those in the unaware group) were able to exploit the cross-trial statistical regularity in the target placement to facilitate their search.

### Positional intertrial priming

Next, we examined for short-term (i.e., inter-trial) positional priming effects (e.g., Allenmark et al. 2019; Sauter et al. 2018; Allenmark, Gokce, et al. 2021) by comparing the mean RTs across the various inter-trial target distances. The results are plotted in Figure 3, where distance 0 means that the target repeated at the exact-same location, which could happen in the random transition condition; distance 1 means that the target moved one position to its previous neighbor, including both the frequent and infrequent directions; all other distances are from trials in the random transition conditions. Positional (inter-trial) priming (e.g., Maljkovic and Nakayama 1996; Krummenacher et al. 2009) would predict an RT advantage for cross-trial repetitions of the target location, providing a strict baseline against which to assess any effect of knowing that the target shifts regularly to the adjacent position in a specific direction across trials.

**Figure 3.**
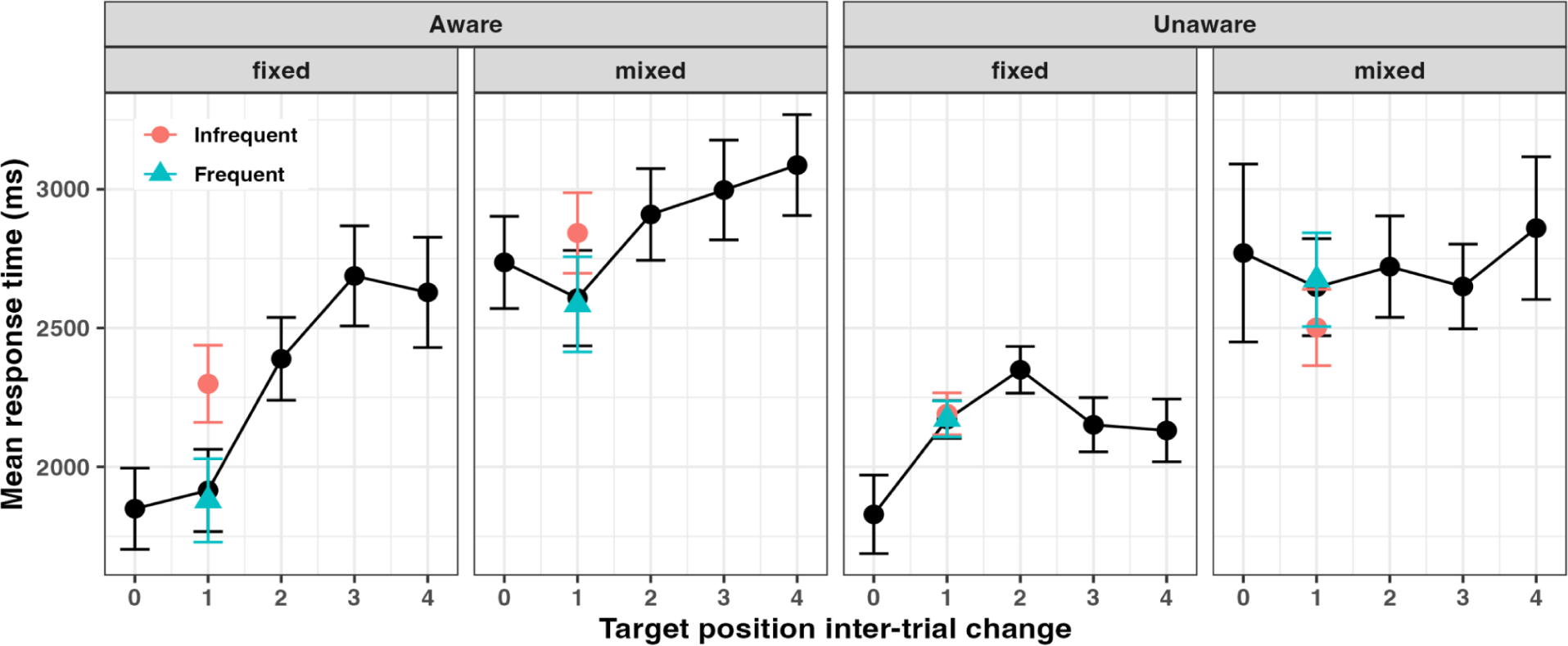
RTs (calculated from individual participants’ medians) as a function of the inter-trial target distance (0 indicates the target repeated at the same location, while 1 denotes the target moved one position to its neighbor, including both the frequent and infrequent directions) in the trial blocks with fixed and mixed target identity, separately for the aware and the unaware groups of participants. Data points marked by green triangles and red circles represent frequent and, respectively, infrequent cross-trial shifts. Error bars represent one standard error of the mean.

An ANOVA of the *distance* effect, with the within-participant factors inter-trial target Distance (0, 1, 2, 3, 4 positions; note that here, distance 1 is the average of shifts in the frequent and the infrequent direction) and cross-trial Target Constancy (target identity fixed, mixed per block) and the between-participant factor Awareness (aware, unaware), revealed the Distance main effect, *F*(4,88) = 9.690, *p* < .001, 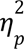 = 0.306, and the Distance × Awareness interaction, *F*(4,88) = 4.813, *p* = .001, 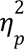 = 0.180, to be significant. Following up the interaction by post-hoc comparisons (with Bonferroni correction) showed that, for the aware group, RTs were significantly faster with both distances 0 and 1 vs. each of the distances 2, 3, and 4, *t*s(23) > 3.87, *p*s < .009, *d*s > 0.592 (there was no difference between distances 0 and 1, *t*(23) = .341, *p* = 1, *d_z_* = 0.052, and among distances 2, 3, and 4, *t*s(23) < 2.268, *p*s > 1, *d_z_*s < 0.346). For the unaware group, by contrast, the distance functions were relatively flat; statistically, there were no significant differences between distances 1, 2, 3, and 4; distance 0 showed some RT advantage (minimum advantage: 101 ms, non-significant; average advantage: 160 ms, *t*(7) = 2.751, *p* = .028, *d_z_* = 0.269). This overall effect pattern was mainly driven by blocks in which the target identity was fixed, which also allowed generally faster search performance.^5^

Thus, there was an advantage for distance 0 – that is, a positional repetition-priming effect – for both the aware and (to a weaker extent) the unaware group, whereas there was an advantage for distance 1 – that is, in this analysis, the combined shift of the target in the frequent and infrequent direction – only for the aware group. This pattern was more prominent in target-fixed blocks of trials, compared to blocks with target identity varying randomly across trials.

Of note, however, for the aware group (and collapsed across the two Target-Constancy conditions), the advantage for distance 1 was entirely due to target shifts in the frequent direction; shifts in the infrequent direction caused a performance slowing relative to both shifts in the frequent direction (infrequent 1 vs. frequent 1, *t*(23) = 3.962, *p* = .004, *d_z_* = 0.598) and exact-same position repetitions (infrequent 1 vs. distance 0, *t*(23) = 3.252, *p* = .033, *d_z_*= 0.491), without a difference between frequent shifts and position repetitions (frequent 1 vs. repetition, *t*(23) = .709, *p* = .48, *d_z_*= 0.107). Again, this pattern was mainly driven by blocks in which the target identity was fixed.

Thus, for the aware group, the positional repetition priming effect was of a comparable magnitude to the dynamic probability-cueing effect. The latter, however, is a genuine effect, rather than simply representing a spatially fuzzy location repetition effect (spreading from the exact same to the neighboring locations), because targets at the location in the infrequent direction (which had the same separation from the 0-distance, reference position as the the frequent location) were associated with a RT cost. Thus, at the very least, one would conclude that the attentional ‘spotlight’ was skewed toward the frequent, and away from the infrequent, direction.

### Inter-trial Priming from Rule-conform (vs. Rule-breaking) Target Shifts

Another possible inter-trial effect might arise from the target on the preceding trial having been positioned consistent with the rule (i.e., having moved to the predicted, frequent location) vs. having shifted in a rule-inconsistent manner (e.g., having moved in the opposite direction to the infrequent location). Rule-consistent shifts might reinforce the rule (or, respectively, inconsistent shifts might weaken the rule), leading to a rule-based inter-trial priming effect. To look for this, we submitted the probability-cueing effect on a given trial *n* to an ANOVA with the within-participant factors Previous (trial *n–1*) Target Location (target at frequent vs. infrequent location) and cross-trial Target Constancy (fixed vs. variable) and the between-participant factor Awareness (aware vs. unaware). The data are plotted in Figure 4.

**Figure 4.**
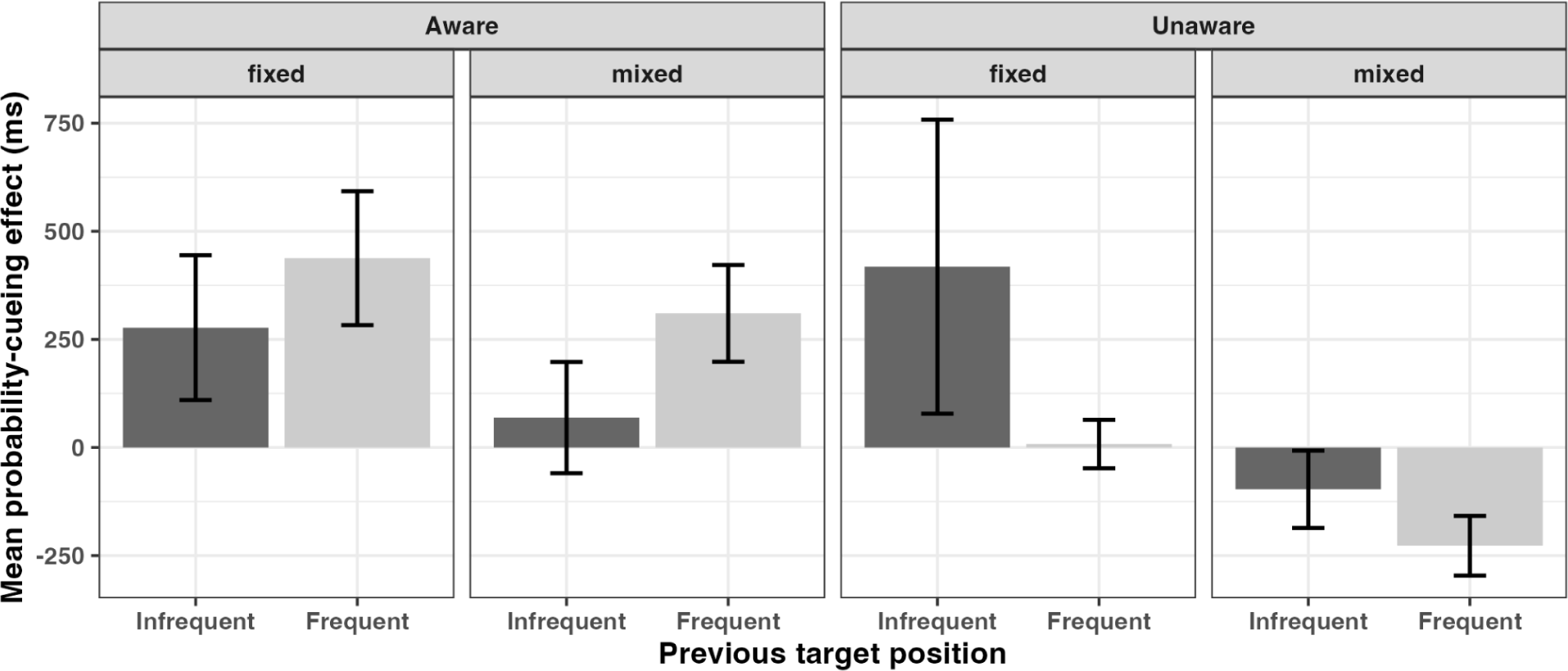
Probability-cueing effect (*RT_infrequent_* − *RT_frequent_*) on a given trial *n* dependent on whether the preceding target (on trial *n–1*) had occurred at the frequent vs. the infrequent location, separately for trial blocks with fixed and mixed target identity and separately for the aware and unaware groups of participants.

There was a main effect of Target Constancy, *F*(1,22) = 5.548, *p* = .028, 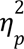 = 0.201, with the probability-cueing effect being greater in target-identity fixed (327 ms) vs. mixed (55 ms) trial blocks. Importantly, the interaction between Previous Target Location and Awareness was significant, *F*(1,22) = 5.944, *p* = .023, 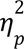 = 0.213, was significant. As can be seen from Figure 4, for the aware group, the probability-cueing effect was numerically greater when the previous target had occurred at the frequent location (i.e., following a rule-consistent shift) compared to an infrequent location (i.e., following a rule-breaking shift), *t*(23) = 1.801, *p* = .513, *d_z_*= 0.364. This pattern appeared to be reversed for the unaware group. In other words, for aware participants, consecutive rule-consistent shifts of the target reinforced the effect of the (discovered) regularity (or, respectively, the effect of the regularity was weakened by a preceding rule-breaking shift). This was not the case for unaware participants, who, by definition, had not discovered the rule.

Of note, in the aware-group, the probability-cueing effect was still significantly positive even when the target appeared at an infrequent location (i.e., after a rule-breaking shift) on the previous trial, *t*(15) = 1.567, *p* = .138, *d_z_* = 0.392. In other words, a rule-breaking shift on the preceding trial just weakened, but did not abolish, the beneficial effect of the regularity.

### Awareness and Dynamic Target-Location Probability Cueing

Figure 5 provides box plots of the probability-cueing (*RT_frequent_* − *RT_frequent_*) effects in the two Target-Constancy conditions, separately for the aware and the unaware groups. An ANOVA of the cueing effect confirmed a significant main effect of the (between-participant) factor Awareness, *F*(1,23) = 8.330, *p* = .009, 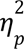 = 0.275: aware participants exhibited an overall greater probability-cueing effect compared to unaware participants (338 ms vs. -76 ms), with the latter actually showing no cueing effect at all (if anything, a negative effect, *t*(7) = 3.011, *p* = .019). Thus, becoming aware of the dynamic, cross-trial regularity in the placement of the target helped participants optimize their search performance.

**Figure 5.**
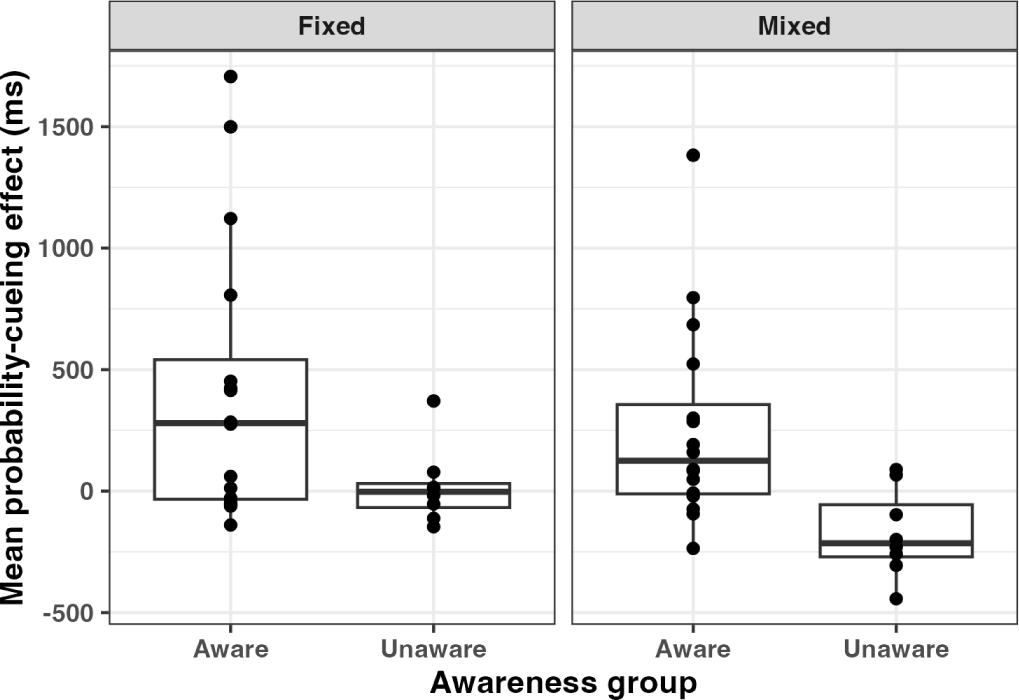
Probability-cueing effect (*RT_infrequent_* − *RT_frequent_*) in the fixed vs. mixed Target-Constancy blocks, separately for the aware and unaware groups of participants.

Next, we calculated the Pearson’s correlation *r* between individuals’ probability-cueing effect and their subjective awareness ratings in response to Q1 (“Did you notice any regularity in the target’s placement across trials?) and, respectively, Q3 (“In what percentage of trials did the target move in the [in Q2 correctly identified/guessed] direction?), separately for the aware and the unaware groups. See Figure 6 for the corresponding plots (scatter plots plus regression lines). In the aware group, a borderline significant correlation was found between for Q1 (*r =* 0.46, *p =* .07 and *R*^2^ *=* 0.21; even with the limited range of confidence ratings: 4,5,6) and a significant correlation for Q3 (*r =* 0.60, *p =* .014 and *R*^2^ *=* 0.36). In contrast and no (or, if anything, numerically negative) correlations were found for unaware participants (Q1: *r =* -0.19, *p =* .65, and *R*^2^ *=* 0.03; Q3: *r = -*0.37, *p =* .35 and *R*^2^ *=* 0.14). Thus, the more confident and the more accurate *aware* participants were in having detected the dynamic cross-trial target-location regularity, the more their search performance benefitted from the regularity – pointing to a critical role of conscious awareness for the manifestation of the dynamic target-location probability-cueing effect.

**Figure 6.**
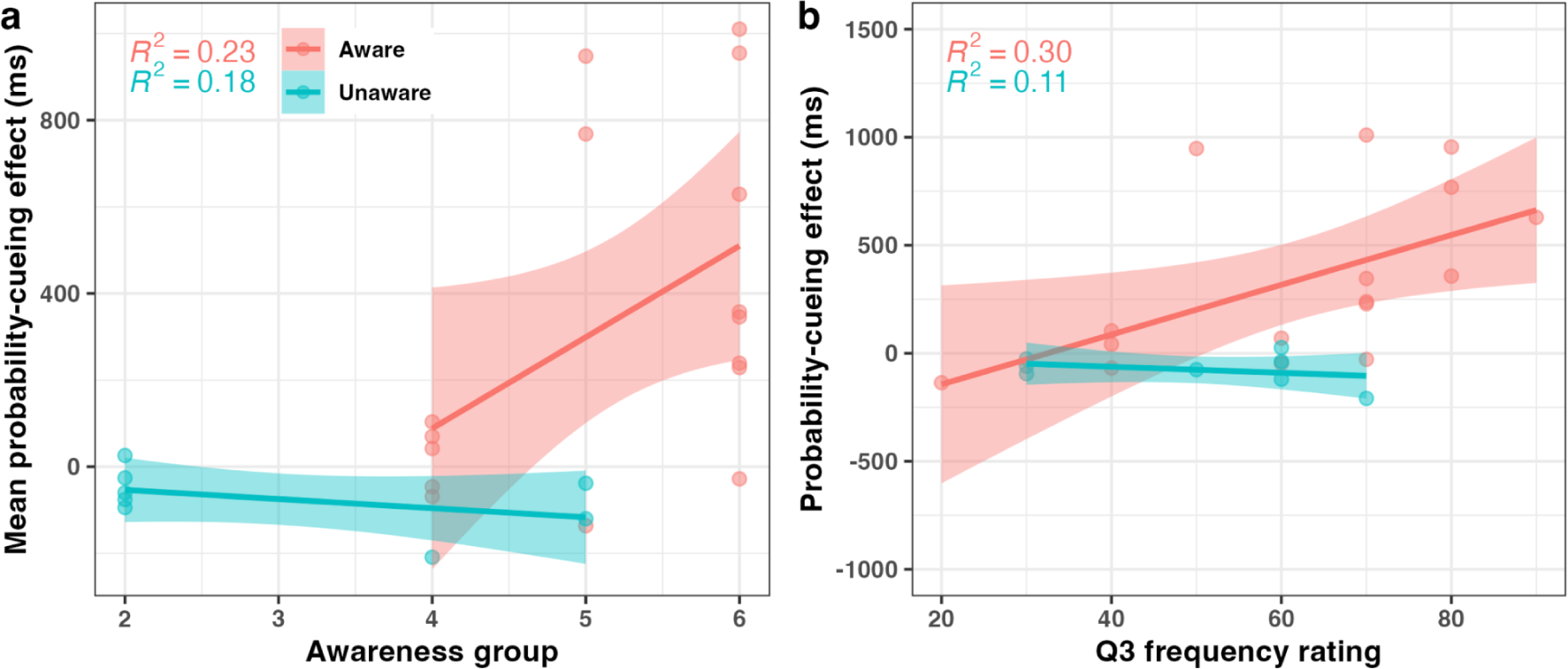
(a) Probability-cueing effect as a function of Q1 confidence rating (1-6), separately for the aware and unaware groups. (b) Probability-cueing effect as a function of Q3 frequency rating (0%–100%). The amount of variance explained by the correlations (r2) between the Q1 and Q3 ratings and the cueing effects are shown at the top of the respective plot. The shaded area represents the 95% confidence interval.

### Eye-movement Results

Due to the absence of a significant probability-cueing effect for the unaware group in the manual RTs, we focused the analysis of the oculomotor behavior on the the aware group (see Appendix B for the results of the unaware group) – aiming to gain a deeper understanding of the underlying mechanisms driving the dynamic probability-cueing effects in a serial search paradigm.

#### Number of saccades until reaching the target and dwell-time on the target

We first examined the average *number of saccades required to reach the target* in an ANOVA with the factors cross-trial Target-Location Transition (frequent, infrequent, random) and cross-trial Target Constancy (fixed, mixed). This ANOVA revealed both main effects to be significant: *F*(2,30) = 13.118, *p* < .001, 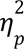 = 0.467 and, respectively, *F*(1,15) = 13.653, *p* = .002, 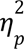 = 0.476. As can be seen from Figure 7a, significantly fewer saccades were required, on average, when the target appeared at the frequent location (4.1 saccades) compared to both the infrequent location (5.5 saccades), *t*(15) = 3.778, *p* = .002, *d_z_* = 1.204, and a random location (5.9 saccades), *t*(15) = 4.885, *p* < .001, *d_z_* = 1.557, without a difference between the later two conditions, *t*(15) = 1.107, *p* = .831, *d_z_* = 0.353. The required number of saccades was also overall lower in fixed target-identity trial blocks compared to randomized blocks, though the difference was not as stark overall (4.9 vs. 5.4 saccades) and of similar magnitude for all Location-Transition conditions (the interaction was non-significant: *F*(2,30) = .048, *p* = .953, 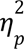 = 0.003).

**Figure 7.**
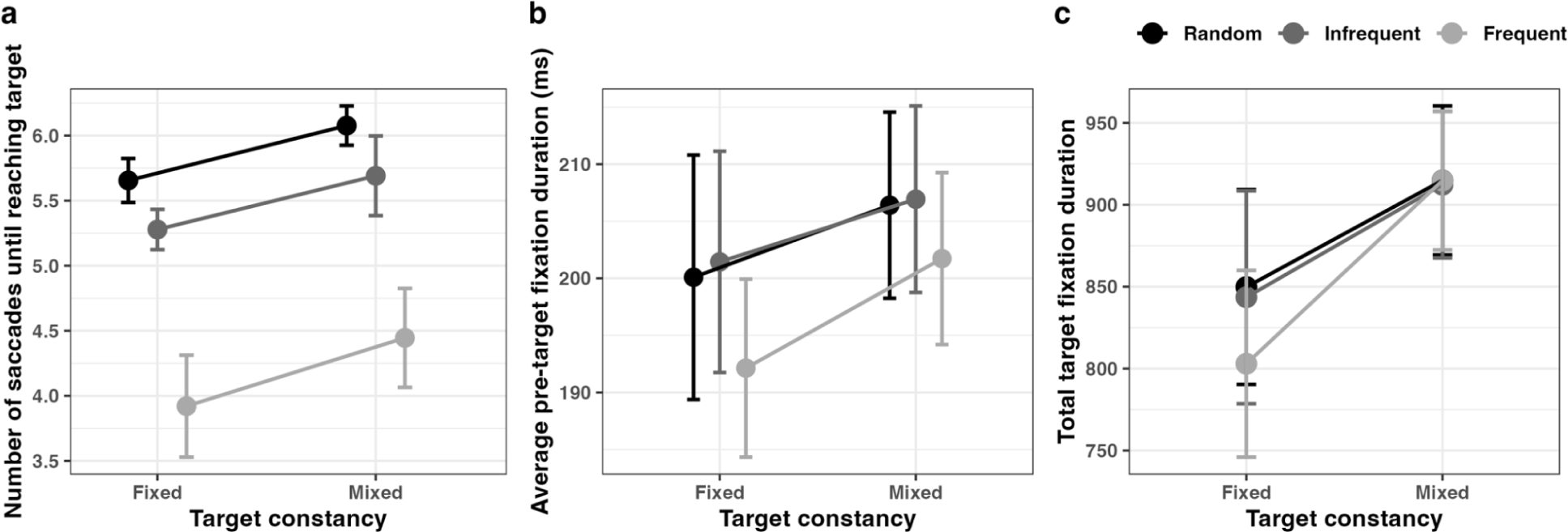
(a) Average number of saccades until reaching the target. (b) Average duration of the fixations before the first saccade to the target, in trial blocks with fixed vs. mixed target identity (cross-trial Target Constancy), dependent on the cross-trial Target-Location Transition (random, infrequent, frequent). Error bars represent one standard error of the mean. (c) Total target fixation duration, in trial blocks with fixed vs. mixed target identity, depending on the cross-trial Target-Location Transition (random, infrequent, frequent). Error bars represent one standard error of the mean.

Thus, the Target-Location effect in the RTs – that is, the expedited RTs to targets at the frequent location – is reflected in the savings in the number of fixational eye-movements required to reach the target positioned at the frequent location.

Figure 7b presents the *average duration of fixations before reaching the target*, in trial blocks with fixed vs. mixed target identity, dependent on the cross-trial Target-Location Transition. A Target-Location Transition × Target-Constancy ANOVA yielded a significant Target-Constancy main effect, *F*(1,15) = 4.638, *p* = .048, 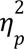 = 0.23: the pre-target fixation durations were by some 10 ms increased in blocks with mixed vs. fixed target identity. They also tended to be reduced for fixations at the frequent vs. the infrequent and random locations, though the Target-Location Transition effect was non-significant (*F*(2,30) = 2.972, *p* = .066, 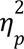 = 0.165) An analogous ANOVA of the *total fixation duration on the target* (see Figure 7c) yielded a significant interaction, *F*(2,30) = 6.448, *p* = .005, 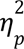 = 0.301 (besides marginal main effects of Target-Location Transition and and Target Constancy, *F*(2,30) = 2.763, *p* = .079, 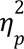 = 0.156, and, respectively, *F*(1,15) = 3.582, *p* = .078, 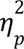 = 0.193). The interaction was due to the fixational dwell-time on the target being shorter in the frequent condition, only in the fixed block (frequent vs. frequent and random combined, 773 ms vs. 828 ms: *t*(15) = 2.812, *p* = .013, *d_z_* = 0.701).

#### First fixation locations

One might assume that participants who learnt the dynamic cross-trial regularity directed their eyes immediately to the frequent target location on a significant proportion of trials. To corroborate this, for the aware group, we analyzed the locations of the very first fixation, that is, the location to which aware participants made the very first saccade on a trial, directly from the central fixation marker. Figure 8c plots the proportions of first fixations directed to the frequent target location, in comparison with the repeated location and the infrequent location, dependent on the target-location cross-trial transition (frequent, infrequent, random), separately for the target-identity fixed and mixed blocks of trials.

**Figure 8.**
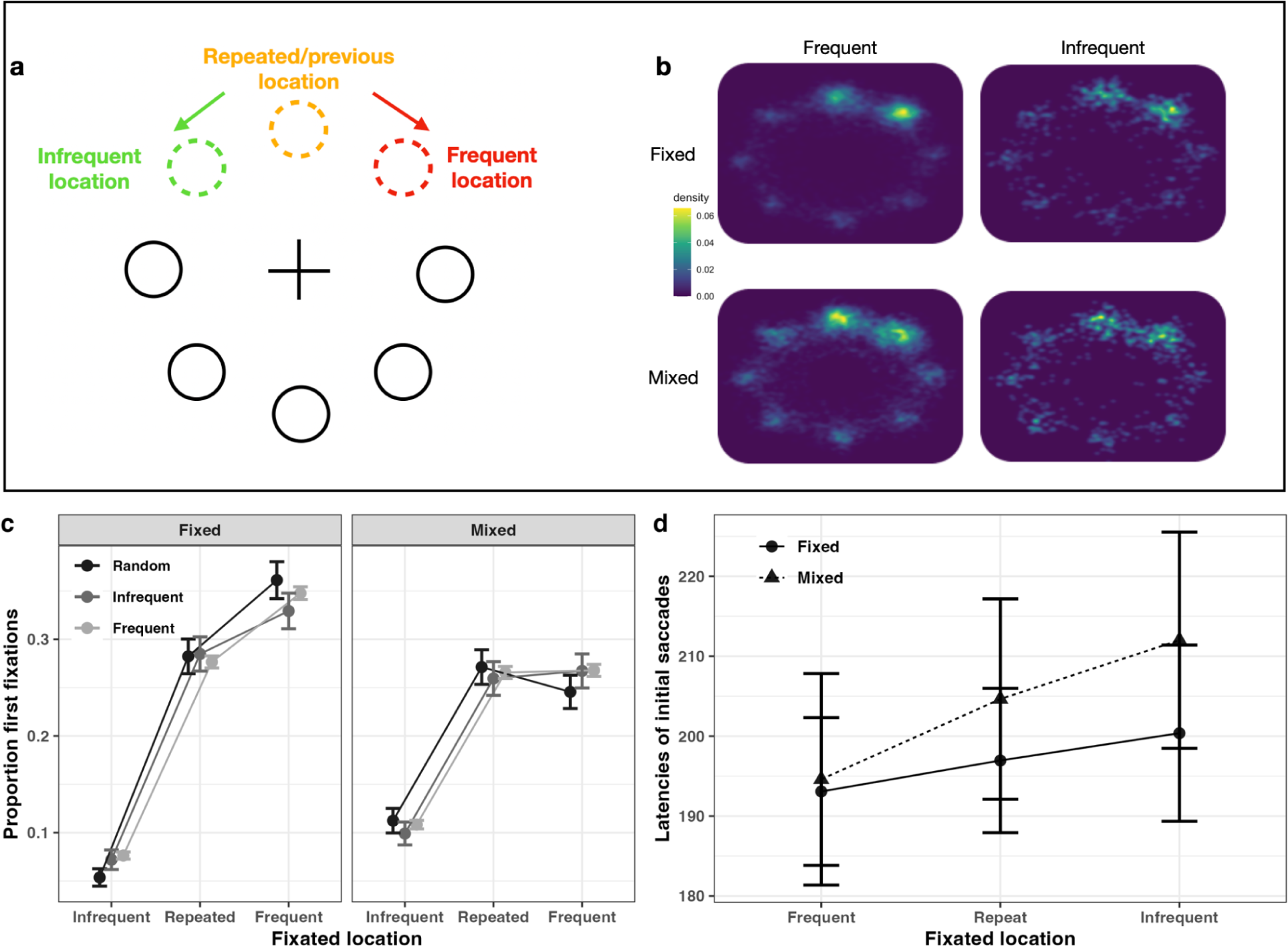
(a) and (b) Heatmaps of the landing positions of the first saccade, depending on the cross-trial Target-Location Transition (frequent, infrequent), for blocks with target identity being fixed vs. mixed (i.e., randomly variable) across trials. As illustrated in 8(a), the fixation locations were rotated such that the target location on trial n-1 is at the top, and the frequent location one to the right, and the infrequent location to the left (for participants with counterclockwise target shifts, the frequent and infrequent locations were flipped right/left flipped). Gaussian filters with smoothing kernels of 0.3° were used to generate all heat maps. (b) Heatmaps for trials on which the target had shifted in the frequent and, respectively, infrequent direction, separately for trial blocks with fixed and mixed target identity. As can be seen, the first saccades were most likely to be directed to the frequent and repeated locations, irrespective of whether the target shifted in the frequent (regular) or the infrequent (irregular) direction; the infrequent location is not more likely to receive a saccade than the random locations (excepting the repeated location). (c) and (d) proportions and, respectively, latencies of initial saccades directed to the frequent, repeated, and infrequent locations (first fixation location) dependent on the cross-trial target-location transition (frequent, infrequent, random), separately for the target-identity fixed and mixed blocks of trials.

A three-way repeated-measures ANOVA of the proportions of first fixation locations revealed a significant main effect of Fixated Location, *F*(2,30) = 5.438, *p* = .010, 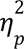 = 0.266. Post-hoc comparisons showed that the frequent location (0.308) was significantly more likely to be the target of the very first saccade than the infrequent location (0.090), *t*(23) = 3.045, *p* = .014, *d_z_* = 1.116; the comparison with the repeated location (0.278) was also significant, *t*(23) = 2.619, *p* = .041, *d_z_* = 0.960, even though the difference was only slight. As can be seen from Figure 8c, this difference derives mainly from the fixed target-identity condition, even though there was no interaction with (or main effect of) cross-trial Target Identity (Fixated-Location × Target-Constancy interaction, *F*(2,30) = 1.619, *p* = .215, 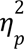 = 0.097).

Of note, the repeated location was prioritized to a similar (though a significantly lesser) degree to the frequent location as the target of the first saccade, reflecting positional intertrial priming. The prioritization of the frequent location, however, is a genuine phenomenon, as the infrequent position (which is equidistant from the repeated location) was clearly deprioritized.

Further of note, there was also no interaction of Fixated Location with the cross-trial Target-Location Transition (*F*(2,30) = 0.049, *p* = .952, 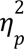 = 0.003). This is interesting because when the first fixation went to the frequent location and the transition was ‘frequent’, the target would actually be located at this position; but when the transition was ‘infrequent’ or ‘random’, the target would not be located at the frequent position. The analogous would apply to the other Fixation-Location conditions. Thus, the lack of a Fixated-Location × Target-Transition interaction implies that the increased proportion of first saccades directed to the frequent location was driven by the discovered regularity; in other words, the rule was applied whether or not the target was actually located there.

An analogous ANOVA of the *latencies of the first saccade* (depicted in Figure 8d) also revealed (only) a main effect of Fixated Location, *F*(2,28) = 3.356, *p* = .049, 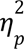 = 0.193. The first saccades were elicited very rapidly upon search display onset, with an average latency of around 200 ms. Post-hoc comparisons revealed the latencies to be significantly shorter for saccades to the frequent vs. the infrequent location (193 ms vs. 206 ms), *t*(14) = 1.939, *p* = .188, *d_z_* = 0.264, with a numerical difference for saccades to the frequent vs. the repeated location (193 ms vs. 201 ms). A distribution analysis revealed the difference between the frequent and infrequent locations to be already evident in the very ‘fastest’ time bins (i.e, the first 22%) of the vicentised latency distributions (*r*^2^(1,5324) = 56.32, *p* < .001), with latencies in the range from between 100 and 150 ms, which would be considered to be too short to be influenced by cognitive control (e.g., Findlay 1997; Sauter et al. 2021).

Interestingly, also, all first saccades in the general direction of the repeated location (i.e., saccades to the frequent, repeated, and infrequent locations) were elicited faster compared to saccades in the other, random directions, the latencies of the latter averaging 220 ms (random vs. frequent: *t*(14) = 4.572, *p* < .001, *d_z_* = 0.582; random vs. repeat: *t*(14) = 3.367, *p* = .010, *d_z_* = 0.428; random vs. infrequent: *t*(14) = 2.433, *p* = .116, *d_z_* = 0.310).

While the landing positions of the first saccades were little influenced by the actual location of the target, a somewhat different picture emerges when looking at the *second* and, especially, the *third* fixation (see Figure 9) in the condition with *fixed* target identity, where the targets located at the frequent location appear to play a role. Examining the cumulative proportions of the first, second, and third fixations falling at a particular location (frequent, repeated, infrequent) as a function of the cross-trial Target-Location transition (frequent, random, infrequent) shows, first, of all, a similar increase in the proportion for the frequent and repeated locations (and a shallower increase for the random locations); that is, both the frequent and the repeated location stay relatively prioritized. Interestingly, though, when the 2nd and, especially, the 3rd fixation fall at the *frequent* location, the cross-trial transition matters: relatively more fixations fall on the frequent location when the target actually occurs there (following a ‘frequent’ transition) compared to when it appears at the infrequent (or a random) location (fixations of *frequent* location: Fixation-Location × Target-Location Transition interaction, *F*(2,30) = 4.77, *p* = .016, 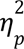 = 0.241; frequent vs. infrequent transition, 3rd fixation: *t*(15) = 4.325, *p* = .002, *d*_z_ = 0.267).

**Figure 9.**
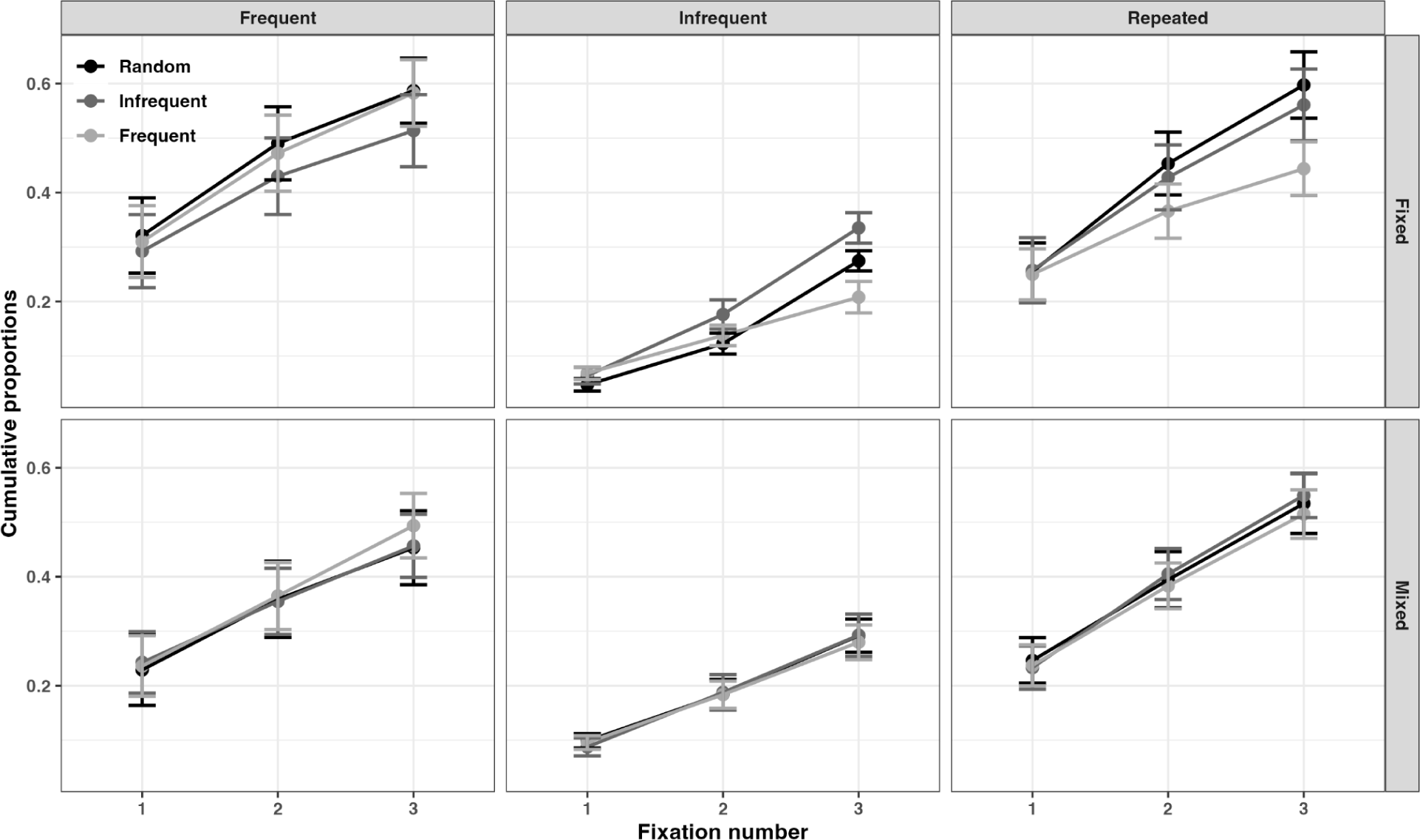
Cumulative probabilities of the first, second, and third fixation falling at a particular location (frequent, repeated, infrequent) as a function of the cross-trial Target-Location transition (frequent, random, infrequent), separately for the fixed and the mixed Target-Identity condition (upper and lower rows, respectively).

Conversely, for the second and, especially, third fixations at the repeated and, respectively, the infrequent location, fewer fixations land at these locations when the target appears at the frequent location (fixations of *repeated* location: Fixation-Location × Target-Location Transition interaction, *F*(2,30) = 3.68, *p* = .037, ^2^ = 0.197; frequent vs. infrequent transition, 3rd fixation: *t*(15) = -3.125, *p* = .055, *d*_z_ = 0.545; 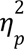 fixations of *infrequent* location: Fixation-Location × Target-Location Transition interaction, *F*(2,30) = 20.03, *p* < .001, ^2^ = 0.572; frequent vs. infrequent transition, 3rd fixation: *t*(15) = -7.630, *p* < .001, *d* = 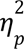 -.1.459). This means that, while the first saccade directed to the frequent location is largely rule-driven, the second and, especially, the third saccade are also influenced by the identity of the item at the frequent location: a target at the frequent location acts like an attractor (over and above the rule-based prioritization of this location), increasing the likelihood of saccades to the frequent location and reducing the likelihood of saccades to random and infrequent locations. This pattern is seen, however, only in the *fixed* Target-Identity condition (in the mixed condition, there was no consistent pattern of interactions), suggesting that it reflects top-down enhancement of critical target features (at the frequent location) by the fixed target template. Interestingly, though, the enhancement appears to be focused on the frequent location.

In the mixed condition, by contrast, the template valid on a given trial can only be established during the search itself – so, there is no (or relatively little) early guidance of search by the target template. This is consistent with an analysis of the saccade patterns ensuing following a first saccade to the target at the frequent location. As depicted in Figure 10, when the target identity is fixed, participants show little tendency to go on to inspect one or two further locations in the immediate neighborhood of the frequent location: in some 50% of the trials, they do not check any location, and in about 25% each they check either one or both neighbors. In the mixed condition, by contrast, they are highly likely to check both neighbors (> 60%) or one neighbor (> 30%) and only very rarely neither (< 10%). This differential pattern (statistically evidenced by a significant interaction between Scanning Pattern [inspection of both, one, or neither neighborhood location] and Target Constancy: *F*(2,30) = 36.159, *p* < .001, 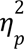 = 0.707, besides a main effect of scanning pattern, *F*(2,30) = 4.599, *p* = .018, 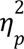 = 0.235) indicates that in the mixed target-identity condition, participants continue scanning to establish the target template valid on a given trial. This would go some way towards explaining why the required number of saccades (and, consequently, the task-final RT) was increased under mixed-identity conditions, as well as why the dynamic cueing effect was somewhat washed out.

**Figure 10.**
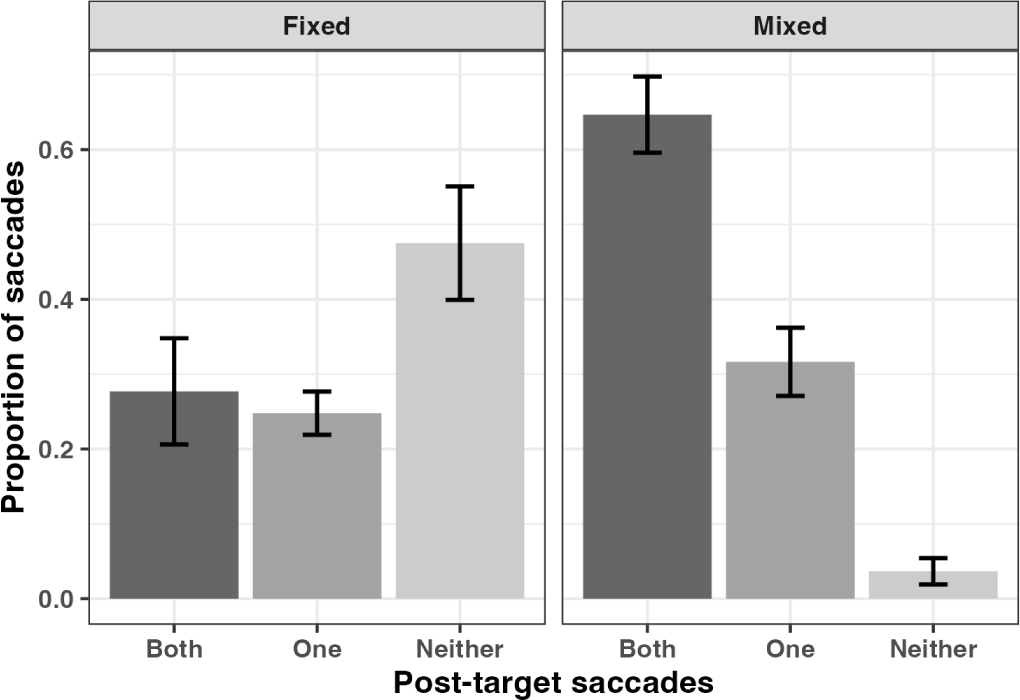
Proportion of saccades directed to one or both neighbors, or neither neighbor, immediately after making the first saccade to the frequent, target-containing location, separately for the fixed and the mixed Target-Identity condition.

#### Awareness and Dynamic Probability Cueing of the First Eye Movement

Consistent with the application of the dynamic rule, the probability-cueing effect in terms of proportions of first saccades directed to the frequent vs. infrequent location correlated marginally with the aware participants’ Q1 confidence rating of the regularity (r = 0.49, *p* = .053 (*r*^2^= 0.24)) and significantly with their Q3 rating of the probability with which the rule applied (r = 0.62, *p* = .01 (*r*^2^= 0.38)) – see Figure 11 for depictions. In other words, the higher aware participants’ estimation of the frequency of, and confidence in, the dynamic regularity, the more likely they were to execute already the first saccade to the frequent, compared to the infrequent, location.

**Figure 11.**
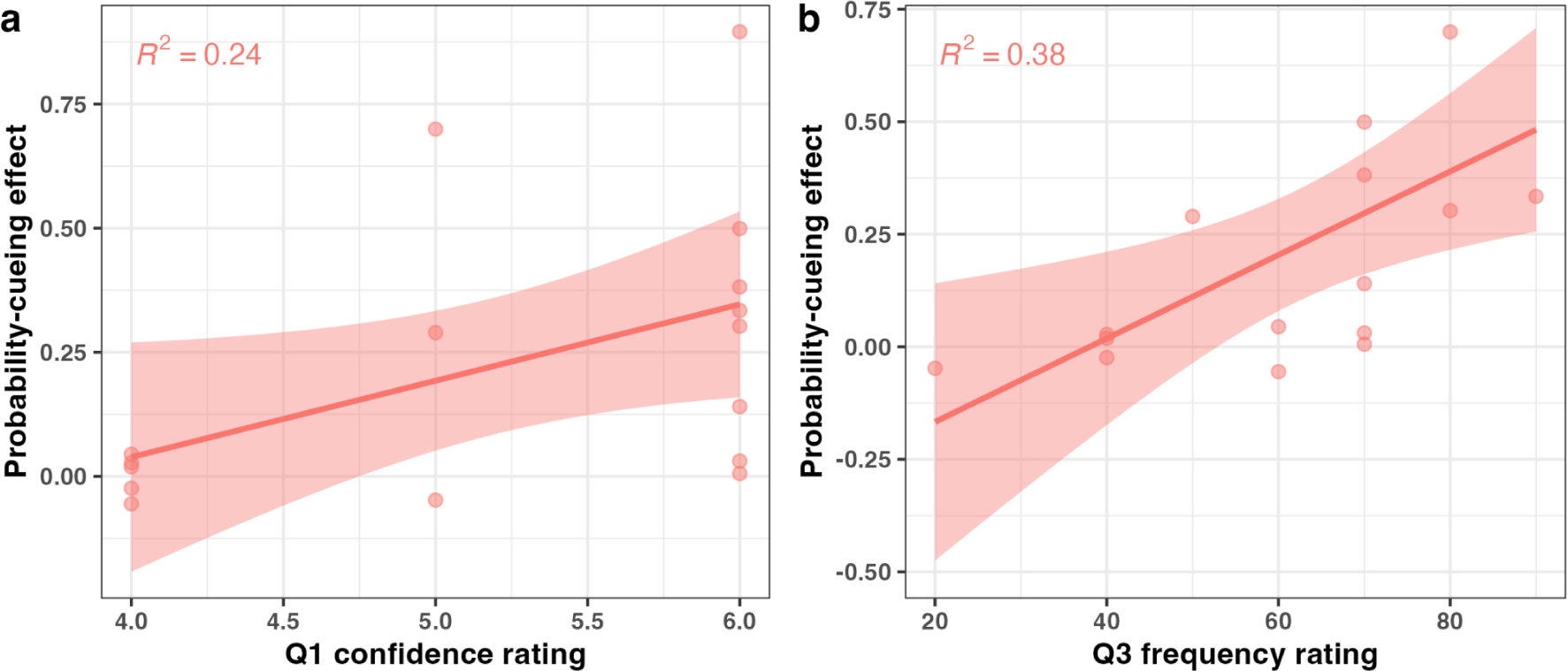
Probability-cueing effect in the first fixation location as a function of the Q1 confidence rating (1-6), for the group of aware participants. (b) Probability-cueing effect in the first fixation location as a function of Q3 frequency rating (0%–100%). The amount of variance explained by the correlations (r2) between the Q1 and Q3 ratings and the cueing effects are also shown at the top of the respective plot.

#### Inter-trial Priming of the First Eye Movement from Rule-conform (vs. Rule-breaking) Target Shifts

Figure 12 provides a plot of the probability-cueing effect in terms of the first eye movement (i.e., proportion of saccades to the frequent minus the infrequent location) dependent on the target location on the previous trial (i.e., trial *n–1* target at frequent vs. infrequent location), separately for trials blocks with fixed vs. mixed target identity. An ANOVA of this cueing effect with the factors Previous (trial *n–1*) Target Location and cross-trial Target Constancy revealed the main effect of Previous Target Location to be significant, *F*(1,15) = 5.255, *p* = .037, 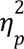 = 0.259: the proportion of first saccades directed to the frequent (vs. the infrequent) location was significantly greater after rule-conforming (.234) than after rule-breaking target shifts (.132) on the preceding trial. Of note, though, the cueing effect was significantly greater than zero even in the latter condition (*t*(15) = 2.378, *p* = .031, *d_z_* = 0.595), consistent with rule violations only weakening, but not abolishing, the effect of the regularity.

**Figure 12.**
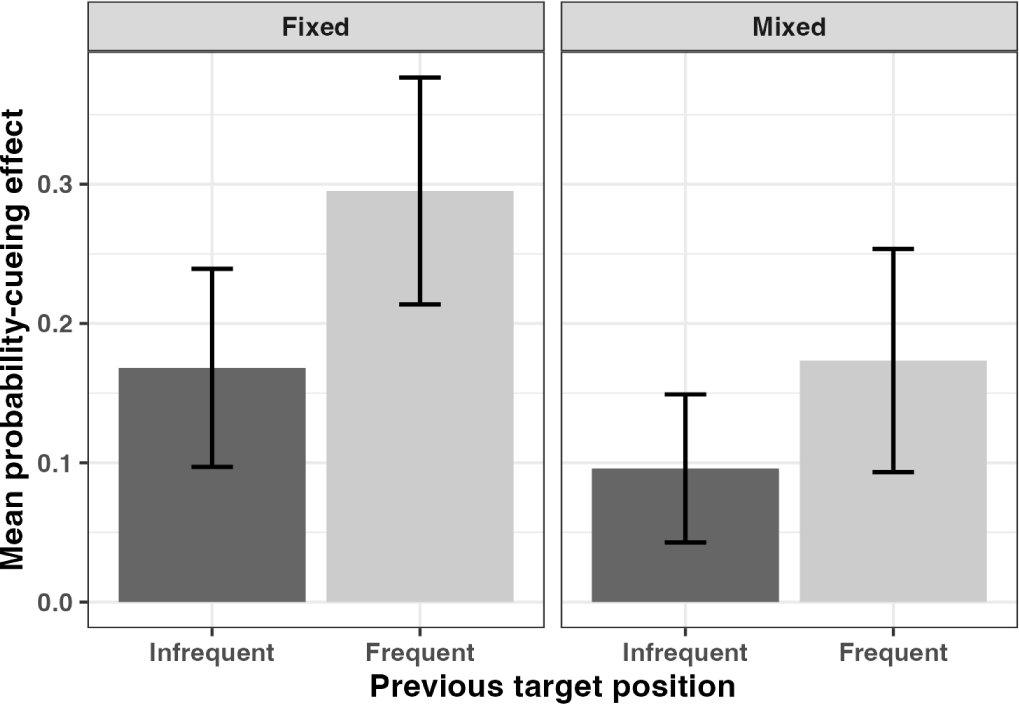
Probability-cueing effect in the first eye movement (proportion of saccades to frequent minus infrequent location) dependent on the target location on the preceding trial (i.e., trial n–1 target at frequent vs. infrequent location), separately for trial blocks with fixed vs. mixed target identity.

## Discussion

The present, eye-tracking study was designed to investigate (1) whether, in a serial search task, participants can learn a dynamic, cross-trial (statistical) regularity regarding the location of the target within a search display and (2), if so, when during the search guidance by the regularity would come into play – the latter by examining (sequential) oculomotor scanning of the displays, as well as the task-final RTs, for evidence of a dynamic target-location probability-cueing effect; further, (3) whether participants’ explicit awareness of the regularity would be systematically related to their probability-cueing effect. In addition to these three main issues, we examined how guidance by the regularity (or rule) compares to positional intertrial priming, how it is modulated by rule-based (i.e., rule-conforming vs. -breaking) intertrial priming, and whether it is influenced by the target identity being known in advance (fixed) vs. having to be established during the task.

The main findings were as follows: Participants (more precisely: ⅔ of participants) successfully learned to utilize the exact-same cross-trial (statistical) regularity in the placement of the target in a serial search task that we had previously shown to be acquired in a parallel, pop-out task (Yu et al. 2023) – a finding that appears to conflict with earlier reports suggesting that the added demands imposed by serial search prevent participants to pick up dynamic regularities (Li and Theeuwes 2020; Li, Bogaerts, and Theeuwes 2022). Importantly, however, of the total sample of participants, only those who (based on a post-experimental awareness test) could be classed as having become aware of the regularity did exhibit a dynamic probability-cueing effect; unaware participants showed no sign of an effect. Of note, in aware participants, search guidance from the discovered regularity kicked in very early: a large proportion of the very first saccades (from the display center) was already directed to the location predicted by the dynamic rule, in a addition to a bias to saccade to the same location that had contained the target on the previous trial; unaware participants displayed only the latter bias. Also, in aware participants, the guidance effect exerted by the dynamic rule was modulated by whether or not the placement of the target on the previous trial was consistent with the rule. Finally, aware participants were able to make use of the rule almost as efficiently when the identity of the target was non-predictable on a trial as when it was fixed. – In the subsequent sections, we consider these findings in more detail.

### Dynamic cross-trial regularities in target placement can be learned even in serial search

The present findings demonstrate that dynamic cross-trial regularities in target placement can be successfully learned and used to optimize performance even in highly demanding, serial search tasks – rather than only in simple pop-out tasks that can be performed spatially in parallel. This at least applies to the regularity implemented here: a shift of the target location, within a circular display arrangement, by one position in either clockwise or counterclockwise direction (fixed per participant) – exactly the same regularity as that used by Yu et al. (2023) in a parallel search task, which allowed for much faster completion times. Interestingly, relative to the random-condition baseline, the performance gains from successfully learning the rule turned out at least as large in the present, serial search task as in the parallel task of Yu et al. (2013): the gains (infrequent minus frequent transition) here amounted to 339 ms, that is, 12.5% of the random-baseline RT (2707 ms) – which compares with a 9.4%-gain (116 ms/1236 ms) in parallel search. In other words, the inherent incentive to acquire the rule was comparable between the two types of task.

Our finding of a cueing effect appears to be at variance with Li and colleagues (Li and Theeuwes 2020; Li, Bogaerts, and Theeuwes 2022), who (using general similar types of display) reported participants to be unable to pick up a different type of dynamic regularity in serial search, which (another sample of) participants could successfully extract in parallel search (learning phase) and subsequently use to expedite serial search (test phase). The main difference between Li and colleagues’ (Li and Theeuwes 2020; Li, Bogaerts, and Theeuwes 2022; Yu et al. 2023) and our studies (Yu et al., 2023; present study) lies in the complexity of the regular cross-trial shift and the frequency with which such shifts were encountered during search. In our design, the proportion of trials on which the target moved to the location predicted by the dynamic regularity (80%) was more than three times larger than that in the design of Li and Theeuwes (only 25%). Also, our dynamic target-location shift was relatively simple: either clockwise or counterclockwise, consistent with how participants might ‘normally’ serially scan a circular search array. The shift introduced by Li and Theeuwes was more complex: if the current target was in, say, the left-most array position, the next target would then invariably appear at the right-most location (but not vice versa). Apart from such shifts occurring only relatively rarely (on some 25% of trials), they would also run counter to normal scanning routines. Thus, it might be that both the frequency with which regular dynamic shifts occur and whether or not they fit with routinized scanning procedures (Seitz et al. 2023) might be critical factors determining whether or not a dynamic regularity is successfully acquired in serial scanning.

Based on the present findings, however, we can conclude that serial search does not per se preclude the possibility that dynamic regularities are extracted and utilized to optimize performance.

### Dynamic target-location probability cueing acts early during search

Over and above the analysis of the task-final RTs, analysis of the oculomotor scanning behavior showed that dynamic target-location probability cueing acts ‘early’ during serial search: already a substantial proportion (some 1/3) of the very first saccades (from the initial fixation marker in the display center) were directed to the predicted, frequent location. Another position receiving almost the same proportion of first saccades was the location that had contained the target on the previous trial, consistent with a positional repetition-priming effect (e.g., Maljkovic and Nakayama 1996; Krummenacher et al. 2009). Of note, at least under conditions with fixed (featural) target identity, a (numerically) greater proportion of first saccades was directed to the predicted vs. the repeated location – that is, the target-location cueing effect tended to dominate the repetition-priming effect. In any case (even under conditions of target-identity swapping), the frequent location received a much greater proportion of first saccades than the infrequent location, even though both were equidistant from the repeated position. This shows that the search priorities (or, respectively, the attentional ‘spotlight’) were systematically biased towards the frequent direction, and away from the infrequent direction. Of note, this early biasing of search turned out to be quite independent of where the target was actually located in the display, that is: it reflects a genuine rule-based effect.

The early prioritization of the frequent and repeated locations was maintained during further scanning, evidenced by these locations continuing to attract the largest proportions of second and third saccades. However, under conditions of fixed target identity, the second and, especially, the third saccade were additionally affected by whether or not the target actually appeared at the predicted, frequent location: a target appearing at the frequent location increased the proportion of (second and third) saccades directed to this location, whereas it decreased the proportions of saccades directed to the repeated and infrequent locations. That is, by the second and third saccade, the priority of the frequent location was not only determined by the dynamic rule; in addition, it became increasingly modulated by the fit of the item at the predicted location to the (fixed) ‘target template’. This suggests that template-based (top-down) enhancement of priority signaling is focused on the predicted location, rather than being ‘broadcast’ equally to all locations (e.g., Wiegand et al., n.d.2023). Interestingly also: the fact that the prioritization of the frequent and repeated locations persisted beyond the first few saccades would imply that the prioritization is coded in scene-based (environmental), rather than retinal, coordinates: the coordinates are dynamically updated across sequential eye movements.

### Rule-based intertrial priming

While the frequent target location is favored as a result of having acquired the dynamic rule, this rule-based prioritization is itself modulated by short-term trial history: it is stronger on a given trial *n* when the shift of the target on the preceding trial *n–1* conformed with the rule (i.e., the target had moved to the then frequent location), or, respectively, it was weaker when the preceding shift violated the rule (i.e., the target had moved to the infrequent location). This effect is seen as early as in the proportions of first saccades, as well as in the task-final RTs. Within a Bayesian framework (e.g., Allenmark, Müller, and Shi 2018; Allenmark, Gokce, et al. 2021), the dynamic rule may be conceived as the acquired long-term ‘prior’ determining the selection priorities, with the weight assigned to the prior on a given trial being modulated by trial history: the current weight is larger following rule-conforming vs. rule-breaking target shifts. Importantly, however, the intertrial weight changes only modulate the effect of the long-term prior, as is evidenced by the fact that there is still significant cueing of the target location following trials on which the rule was violated; that is, the weight assigned to the prior is not set to zero.

To our knowledge, this rule-based intertrial priming effect is a novel phenomenon that has not been reported before. Of course, there are reports of intertrial priming effects associated with statistical learning of static regularities. For instance, the interference caused by a salient distractor is increased when the distractor occurs at a previous target location, while it is reduced when it occurs at a previous distractor location; conversely, search is expedited when the target occurs at a previous target location, while it is slowed when it occurs a previous distractor location (see, e.g., Sauter et al. 2018), and this may be modulated by a static ‘rule’, that is, how likely the target or, respectively, a distractor, occurs at a particular, fixed location. However, these are essentially positional intertrial effects, attributable to some facilitatory or inhibitory ‘tags’ having been placed on the respective position as a result of having encountered a target or a distractor there on trial *n–1*. In the present, dynamic scenario, however, there was, by definition, never a target at the predicted location on trial *n–1*, arguing that the priming effect is genuinely rule-related. Nevertheless, of course, it may exert its effect in location-based coordinates, such as on a common map representing attentional (and oculomotor) priorities.

### Dynamic probability-cueing is modulated but not abolished by target-identity swapping

Further of interest, dynamic target-location probability cueing was not abolished by random swapping of the target identity across trials. However, under conditions of swapping, the search RTs were overall prolonged (associated with an increased number of fixations) and the cueing effect was reduced, from 420 ms in fixed- to 257 ms in mixed-identity blocks in the aware group. This is not surprising as overall, and especially on identity-swap as compared to -repeat trials, more fixations were necessary to establish exactly what was the searched-for target and what the irrelevant non-targets on a given trial. Thus, even when the first saccade was directed to the predicted location, further steps of processing – involving comparisons with (and saccades to) the neighboring items – would have been necessary to ascertain the target identity. This is exacerbated on identity-swap trials, where the ‘default’ assumption that the target (identity) stays the same as on the previous trial is proved to be wrong and the ‘target template’ has to be changed – as evidenced by a target-identity- (or ‘template’-) based repetition-benefit/change-cost effect (mirroring feature-based priming effects in pop-out or, respectively, feature-conjunction search; (e.g., Maljkovic and Nakayama 1994; Kristjánsson, Wang, and Nakayama 2002; Geyer, Müller, and Krummenacher 2006). Interestingly, however, even though the probability-cueing effect was reduced on identity-swap (vs. -repeat) trials, it remained significantly larger than zero. In other words, having acquired the dynamic regularity in the positioning of the target across trials did facilitate performance even under the most demanding search conditions.

Whether these conditions allow the dynamic regularity to be efficiently acquired in the first instance is a different question. Our data are non-conclusive in this regard. For the first four (of the total eight) blocks, the cueing effect differed little between aware participants starting with the fixed vs. those starting with the mixed target-identity condition; the latter group, however, tended to show an increased (numerically nearly doubled) effect after the switch to the fixed condition, while the former did did not exhibit any gain following the switch to the mixed condition. While the critical interaction was non-significant (*F*(1,14)^6^ = 0.86, *p* = .369), this pattern is more consistent with the mixed target-identity condition interfering with the ‘expression’ of the cueing effect, rather than impeding the acquisition of the dynamic regularity as such. The expression of the effect would be affected due to the need to establish the target template valid on a trial even if the target at the frequent location target was the first item inspected (see above).

### Awareness of the dynamic rule and target-location probability cueing in serial search

The last, but not least important, point of discussion concerns the role of awareness for the dynamic target-location probability-cueing effect in serial search. In contrast to the majority of studies of probability-cueing effects, which concluded that spatial statistical learning is not dependent on awareness and thus ‘implicit’ in nature (e.g., Jiang, Swallow, and Rosenbaum 2013; Jiang, Won, and Swallow 2014; Won and Jiang 2015), we found strong evidence of the present, dynamic target-location cueing effect involving awareness. First of all, only participants classed as ‘aware’ (⅔ of participants) based on our post-experimental questionnaire showed a dynamic cueing effect in both the task-final RTs and the earliest eye movements; ‘unaware’ participants (⅓), by contrast showed no sign of a cueing effect in either early or late(r) measures of performance (they only exhibited a tendency to saccade to the previous target location). Second, in ‘aware’ participants, the strength of the cueing effect, even in the proportion of first eye movements directed to the predicted location, correlated significantly with how realistically they believed the rule applied: the more accurately participants estimated the frequency with which the target shifted in the regular direction, the larger their cueing effect. Showing an element of ‘explicitness’ is in line with other studies that used more sensitive awareness tests (e.g., Giménez-Fernández et al. 2020; Golan and Lamy 2023), as well as our own experiment of dynamic target-location cueing in *parallel* search (Yu et al. 2023). In particular, it is in line with the significant correlation reported by Giménez-Fernández et al. (2020), whose measures of awareness we adopted in present study. Interestingly, however, here a role of awareness was seen in a relatively small sample (in 16 out of a total of 24 participants) – arguing that, at least in the present, dynamic scenario, a large sample size may not be crucial for demonstrating ‘awareness’.

However, exactly what is the role of awareness in the dynamic cueing effect? The effect is, in some way, dependent on awareness, as only the group of ‘aware’ participants showed a benefit, but not the ‘unaware’ group. But, despite a significant correlation between awareness of the dynamic regularity and the cueing effect, does this mean that this effect is a ‘voluntary’ in nature, that is, mediated by participants deliberately applying the rule to guide their search on each (or most) trial(s)? While this is a possibility, recall that the latencies of first saccade to the predicted location were rather short (some 190 ms), as were, in fact, the latencies to the repeated and infrequent locations (somewhat over 200 ms) which were both shorter compared to random locations (> 220 ms). This pattern suggests an ensuing competition, upon display onset, of the search items at locations in the general direction of the previous target position (to which the task had just required a saccade to be executed), for which activity remains elevated across trials on some (integrative) oculomotor priority map, likely, in the superior colliculus (e.g., Veale, Hafed, and Yoshida 2017). Thus, while the repeated location remains a strong attractor for the next eye movement (the first saccade on the new trial), this competition is then resolved in favor of the frequent location, perhaps through a rule-related input injected into the priority representation via frontal-eye-field neurons that represent the dynamically updated, ‘goal’-related priority. Given that the display array was not visible during the intertrial interval (there were no placeholders), the updating of the saccade goal may happen only after search-display onset. In this case, latencies (well) below 200 ms may not be sufficient for consciously mediated inputs to influence saccade programing.^7^ Accordingly, one would have to assume that rule-based dynamic goal updating, while perhaps initially requiring conscious control to be set up, eventually becomes a rather automatized, ‘implicit’ process that runs off without ‘explicit’ cognitive intervention (cf. Shiffrin and Schneider 1977). Thus, it may be premature to conclude from the correlation between awareness of the dynamic regularity and the cueing effect that this effect is causally mediated by awareness on each (or most) trial(s).

Overall, it remains that there is no *dynamic target*-location probability-cueing effect in *serial* search when there is no awareness of the regularity. In Yu et al. (2023), where we implemented the same cross-trial regularity, we argued that this also applies to dynamic *target*-location cueing in *parallel* search. By implication, we attributed our finding that the same regularity did *not* produce a cueing effect when it was implemented in a pop-out *distractor* in *parallel* search to the fact that participants did *not* become of the regularity in the cross-trial distractor-location shift – whereas participants became aware of the exact-same shift when implemented in the pop-out target.^8^ Thus, we propose that participants becoming aware of the regularity (and, on the part of the experimenter, establishing awareness by sensitive measures; cf. Vadillo, Konstantinidis, and Shanks 2016; Vadillo et al. 2020; Giménez-Fernández et al. 2020) is crucial for dynamic probability-cueing effects in any type – serial or parallel – of search to develop.

## Conclusion

Our findings show that, contrary to previous reports, participants *can* extract dynamic regularities in the cross-trial placement of the target even in *serial* search (involving sequential eye movements) and utilize them to improve task performance – at least when the regular cross-trial target shift is relatively simple and occurring frequently. This finding is non-trivial, as the exact-same regularity is not picked up when implemented in a salient, ‘pop-out’ distractor in parallel search (Yu et al. 2023). Crucially, this dynamic target-location probability-cueing effect is evident even in (both the proportion and latency of) the very first saccade elicited upon search-display onset, which is purely driven by the learnt rule and not the actual location of the target in the display. Further, it correlates with participants’ awareness of the dynamic regularity. However, given how fast the rule-injected bias can operate after display onset (it is evident already in the very fastest first saccades, elicited between 100 and 150 ms post-display onset), the cueing effect itself may not be consciously mediated. In this case, though, awareness would play a crucial role in acquiring the effect in the first instance. Alternatively, the rule-based biasing may already be ‘prepared’ in the intertrial interval, allowing the cueing effect to ramp up rapidly after search display onset. More work, for instance involving electrophysiological measures, is necessary to clarify this. Also, further work would be required to map the boundary conditions, in terms of both the complexity of dynamic target regularities and the frequency with which they occur, for a cueing effect to be observable.

## Supplementary

### Appendix A: extra behavioral analysis

#### Does target-identity swapping influence performance in mixed trial blocks?

To check whether target swapping might affect performance in mixed-block trials, we conducted a 2×2×3 ANOVA with the within-participant factors cross-trial Target Identity (repetition vs. switch), and cross-trial Target-Location Transition (frequent, infrequent), and the between-participant Awareness (aware vs. unaware). This ANOVA revealed RTs to be overall faster on trials in which the target was repeated rather than switched, *F*(1,22) = 10.538, *p* = .004, 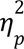 = 0.324 (Target-Identity main effect). There was also a marginal interaction between Awareness and the Target-Location Transition, due to the aware (but not the unaware) participants exhibiting a probability-cueing effect (i.e., faster RTs to targets at frequent vs. infrequent locations). See Figure A1 for a depiction of this effect pattern.

**Figure A1.**
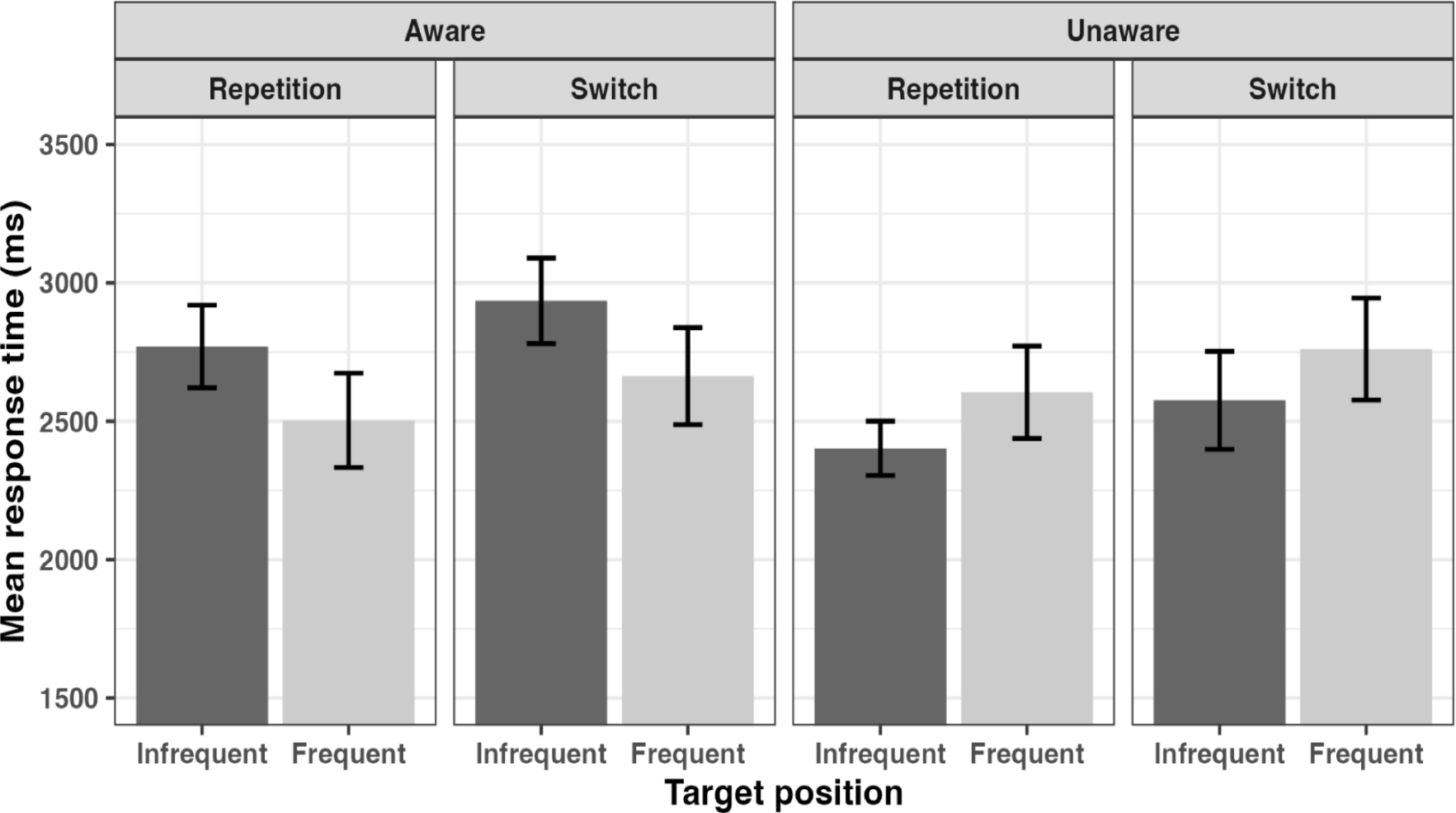
RTs as a function of the cross-trial target-location transition (the frequent, infrequent) for trials *n* in which the target (identity) repeated vs. switched relative to trial *n–1*, separately for the aware and unaware groups of participants. Error bars represent one standard error.

#### Appendix B: Analysis of Eye-movements in the Unaware Group

Figure B1(a) presents the mean number of saccades required to reach the target, in trial blocks with fixed vs. mixed (i.e., randomly varying) target identity, for the three cross-trial target-location transition conditions (frequent, infrequent, random). An ANOVA revealed the Target-Constancy × Target-location Transition interaction to be significant, *F*(2,14) = 5.023, *p* = .023, 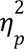 = 0.418. Although post-hoc tests yielded no significant comparisons, the interaction appears driven by the increased saccade numbers in the frequent and random transition conditions vs. the infrequent condition in mixed trials blocks. Importantly, there was no systematic advantage for frequent vs. infrequent and random transitions in either (fixed, mixed) type of block.

**Figure B1.**
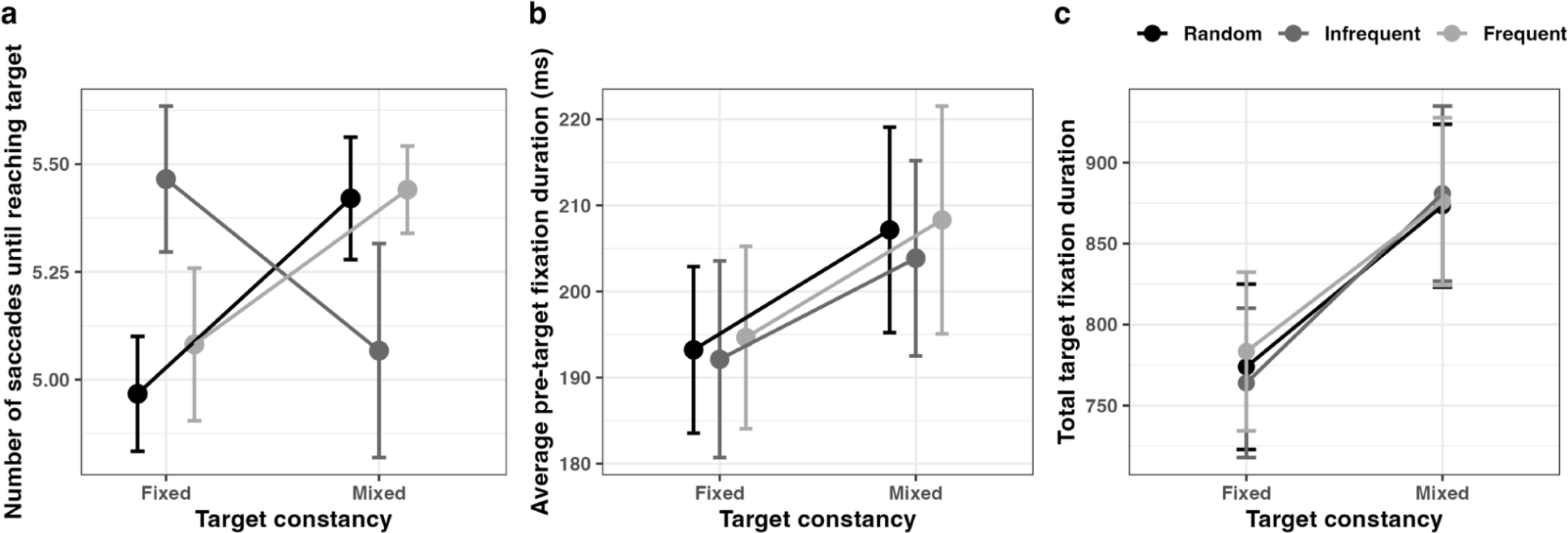
(**a**) Number of saccades until reaching the target under conditions of fixed vs. mixed target-identities, for the three cross-trial target-location transition conditions (frequent, infrequent, random). Error bars represent one standard error of the mean. (**b**) Average pre-target fixation duration, in trial blocks with fixed vs. mixed target identity (cross-trial Target Constancy), dependent on the cross-trial Target-Location Transition (random, infrequent, frequent). Error bars represent one standard error of the mean. (**c**) . Total target fixation duration, in trial blocks with fixed vs. mixed target identity (cross-trial Target Constancy), dependent on the cross-trial Target-Location Transition (random, infrequent, frequent)for target transition conditions (the random, infrequent, and frequent). Error bars represent one standard error of the mean

Figure B1(b) presents *average pre-target fixation duration*, in trial blocks with fixed vs. mixed target identity (cross-trial Target Constancy), dependent on the cross-trial Target-Location Transition (random, infrequent, frequent) for the unaware group. A repeated-measures ANOVA of the *average pre-target fixation duration*, with the factors cross-trial Target-Location Transition (frequent, infrequent, random) and Target Constancy (mixed vs. fixed), yielded only a main effect of Target Constancy, *F*(1,7) = 6.525, *p* = .038, 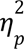 = 0.482: pre-target fixations were shorter in blocks with fixed vs. randomly varying target identity (193 ms vs. 206 ms).

Figure B1(c) presents the total target fixation duration in trial blocks with fixed vs. mixed target identity, for the three cross-trial target-location transition conditions (frequent, infrequent, random). An ANOVA yielded only a main effect of the Target Constancy: *F*(1,7) = 6.937, *p* = .034, 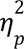 = 0.498, with the fixation duration on the target being shorter in blocks with fixed vs. randomly varying target identity (739 ms vs. 844 ms).

**Figure B5.**
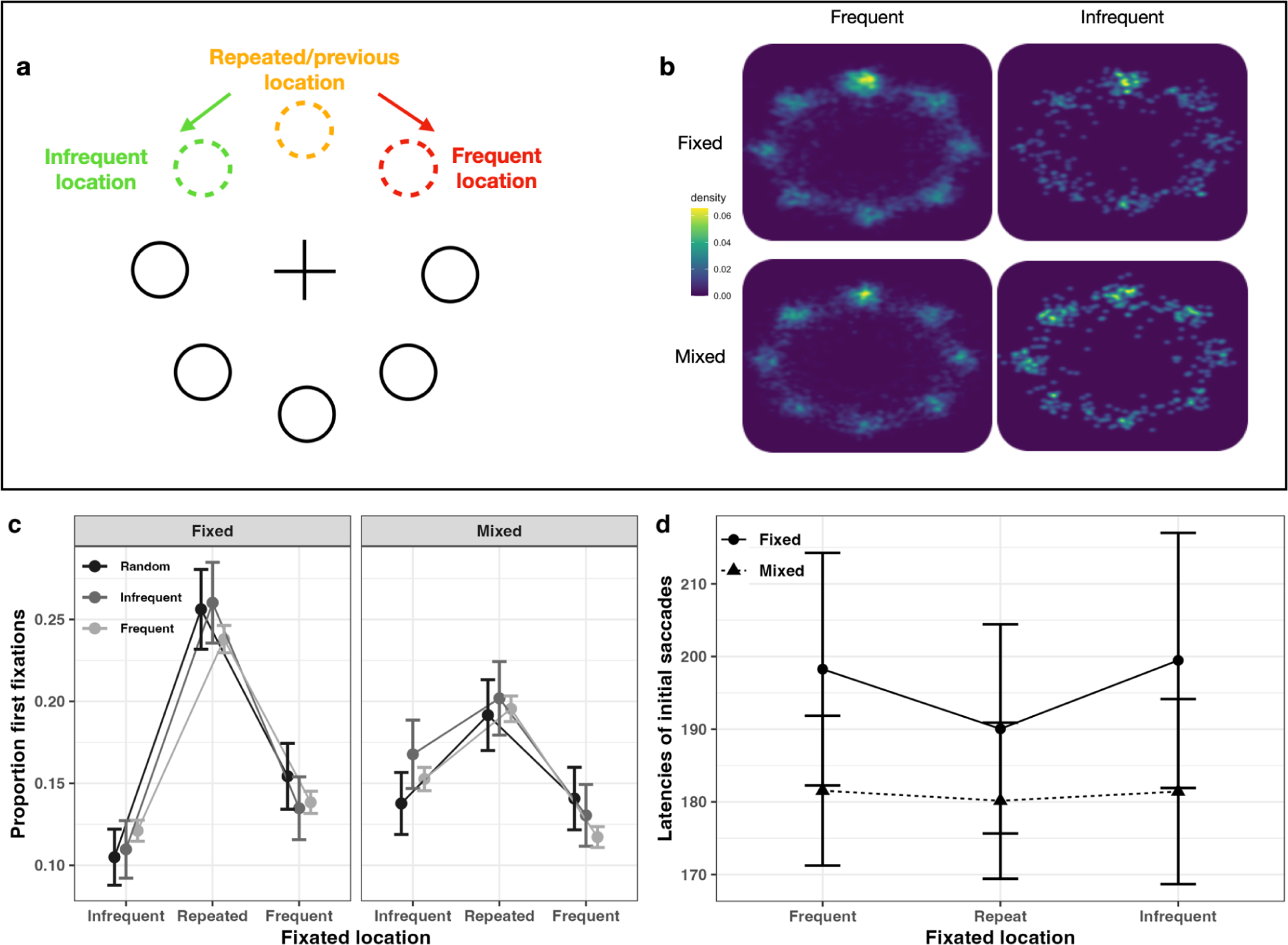
**(a**) and (**b**) Heatmaps of the landing positions of the first saccade, depending on the cross-trial Target-Location Transition (frequent, infrequent), for blocks with target identity being fixed vs. mixed (i.e., randomly variable) across trials. As illustrated in 8a, the fixation locations were rotated such that the target location on trial *n-1* is at the top, and the frequent location one to the right, and the infrequent location to the left (for participants with counterclockwise target shifts, the frequent and infrequent locations were flipped right/left flipped). Gaussian filters with smoothing kernels of 0.3° were used to generate all heat maps. **(b)** Heatmaps for trials on which the target had shifted in the frequent and, respectively, infrequent direction, separately for trial blocks with fixed and mixed target identity. As can be seen, the first saccades were most likely to be directed to the repeated location; the frequent location was *not* more likely to receive a saccade than the infrequent or random locations (excepting the repeated location). (**c**) and (**d**) proportions and, respectively, latencies of initial saccades directed to the frequent, repeated, and infrequent locations (first fixation location) dependent on the cross-trial target-location transition (frequent, infrequent, repeated), separately for the target-identity fixed and mixed blocks of trials.

For the group of unaware participants, there were no significant correlations between the probability-cueing effect in the first fixations and the confidence they associated with their Q1 response (*r* = 0.42, *p* = .31 (*r*^2^ = 0.19) and the rated frequency in their Q3 response (*r* = –0.37, *p* = .35 (*r*^2^ < 0.01))

**Figure B6.**
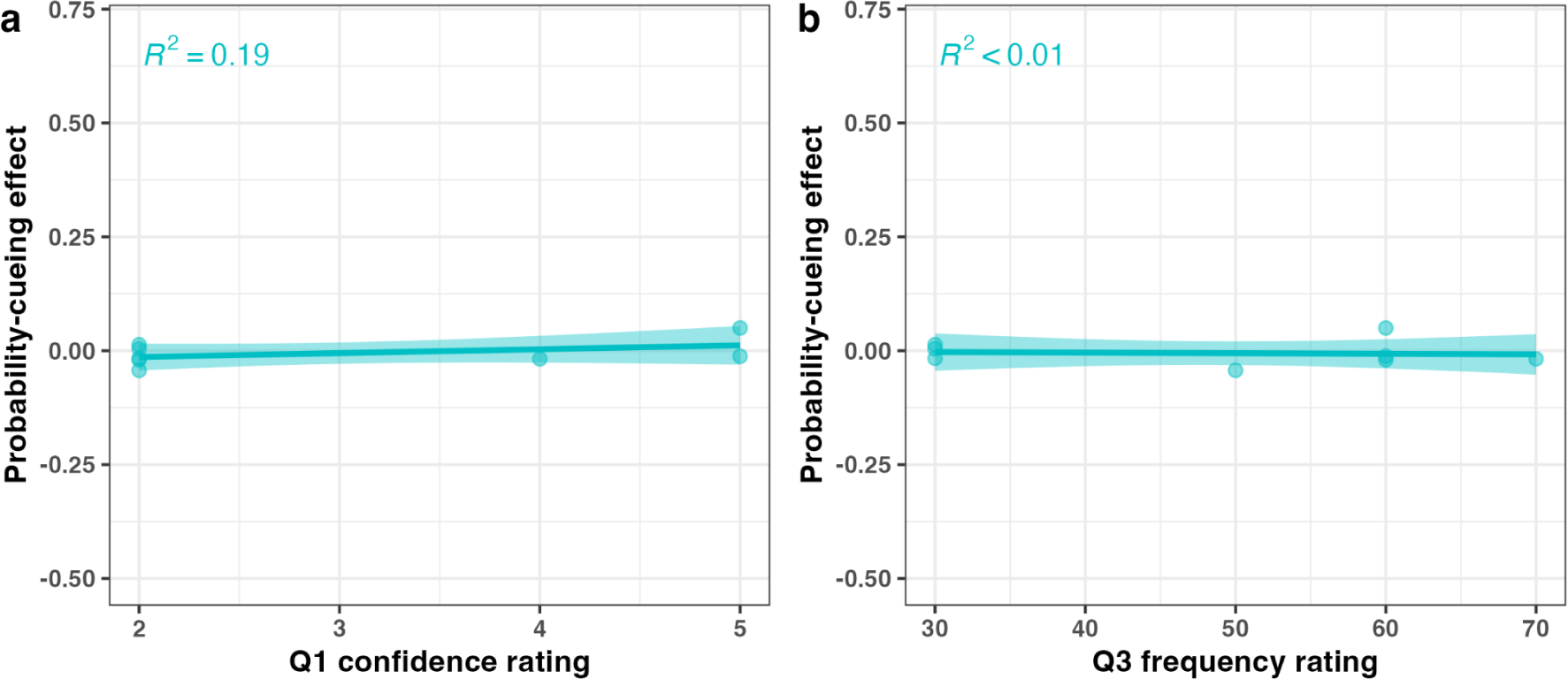
Probability-cueing effect in the first fixation location as a function of the Q1 confidence rating (1-6), for the group of unaware participants. (**b**) Probability-cueing effect in the first fixation location as a function of Q3 frequency rating (0%–100%). The amount of variance explained by the correlations (*r*^2^) between the Q1 and Q3 ratings and the cueing effects are also shown at the top of the respective plot.

#### Inter-trial Priming of the First Eye Movement from Rule-conform (vs. Rule-breaking) Target Shifts (for unaware group)

Figure B7 provides a plot of the probability-cueing effect in terms of the first eye movement (i.e., proportion of saccades to the frequent minus the infrequent location) dependent on the target location on the previous trial (i.e., trial *n–1* target at frequent vs. infrequent location), separately for trials blocks with fixed vs. mixed target identity. An ANOVA of this cueing effect with the factors Previous (trial *n–1*) Target Location and cross-trial Target Constancy revealed no significant effects. In particular, the main effect of Previous Target Location was non-significant, *F*(1,7) = 0.114, *p* = .746, 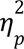 = 0.016), that is, the proportion of first saccades directed to the frequent (vs. the infrequent) location was not different following rule-conforming (-0.003) as compared to rule-breaking target shifts (-0.01) on the preceding trial.

**Figure B7.**
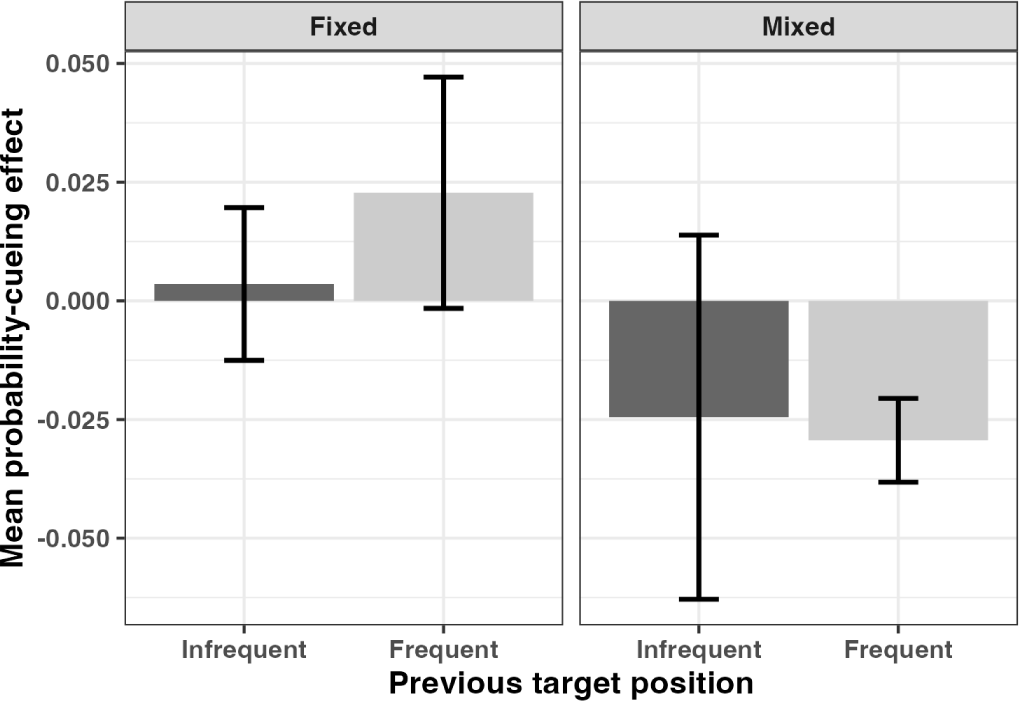
Probability-cueing effect in the first eye movement (proportion of saccades to frequent minus infrequent location) dependent on the target location on the preceding trial (i.e., trial *n–1* target at frequent vs. infrequent location), separately for trial blocks with fixed vs. mixed target identity.

## Footnotes

1 In Li and Theeuwes’s (2020) design, regular target shifts occurred in only 25% of the trials, and, following a target event at a critical – say, the left-most – position, the next target would be appear invariably (i.e., with 100% probability) at the right-most position; so, the rule was deterministic (rather than being probabilistic), and ‘predicted’ position was fixed (rather dynamically changing). In Yu et al. (2023), by contrast, regular target shifts – by one position in, say, clockwise direction – occurred in 80% of the trials; so, the rule was probabilistic and predicted position changed systematically.

2 Li et al. (2022) implemented only a serial search condition. They found that when the target was purely shape-defined throughout the experiment, there was no dynamic target-location probability cueing effect. But when the target was made a color singleton (pop-out) item during an initial learning phase, participants acquired a cueing effect, which persisted in a subsequent test phase in which the color information was removed.

3 This would also be consistent with Li et al. (2022), where only two of a total of 57 participants could be said to have become explicitly aware of the dynamic regularity implemented in their study: failure to become aware of the regularity would predict the absence of a cueing effect.

4 Some eye-tracking studies investigating *static* probability-cueing effects found more first saccades to be directed to high-vs. low-probability target regions and reported that hardly any participants became aware of the static regularity (Jiang, Won, and Swallow 2014; Addleman and Lee 2022). However, only a few pertinent studies thoroughly assessed participants’ level of awareness of the manipulated probabilities(Vicente-Conesa et al. 2021; Giménez-Fernández et al. 2020; Yu et al. 2023; Golan and Lamy 2023).

5 In more detail, the Distance effect was also modulated by Target Constancy (Distance x Constancy interaction, *F*(1,22) = 6.925, *p* < .001, ηp2 = 0.239), in addition to RTs being overall faster in target-fixed vs. mixed trial blocks (main effect of Target Constancy, *F*(1,22) = 44.123, *p* < .001, ηp2 = 0.667). For target-fixed blocks, RTs differed significantly between 0, 1 vs. 2, 3, and 4 (*t*s(23) > 3.395, *p*s < .039, *d*s > 0.542), except distance 0 vs. 1 (*t*(23) = 2.118, *p* = 1.0, *d_z_*= 0.338). For mixed blocks, RTs did not differ among any of the distances (apart from a small difference between distances 1 and 4, *t*(23) = 3.596, *p* = .02, *d_z_* = 0.574).

6 The degrees of freedom are reduced because one of the aware participants had insufficient trials in one of the conditions and was so excluded from analysis.

7 This would also be consistent with Findlay (1997), who concluded from his study of saccade target selection during pop-out and feature-conjunction searches that “the generation of the first saccade is a relatively automatic process, rather than one which is subject to cognitive control” (p. 628).x

8 This would also explain Li and Theeuwes’ (2020) non-finding: their participants did not become unaware of their (more complex and less likely) dynamic target-location regularity and accordingly exhibited no cueing effect.

